# A system-view of *B. pertussis* booster vaccine responses in adults primed with whole-cell vs. acellular vaccine in infancy

**DOI:** 10.1101/2020.05.15.098830

**Authors:** Ricardo da Silva Antunes, Mikhail Pomaznoy, Ferran Soldevila, Mariana Babor, Jason Bennett, Yuan Tian, Natalie Khalil, Yu Qian, Aishwarya Mandava, Richard H. Scheuermann, Mario Cortese, Bali Pulendran, Christopher D. Petro, Adrienne Gilkes, Lisa A. Purcell, Alessandro Sette, Bjoern Peters

## Abstract

Whole-cell inactivated vaccine against *Bordetella pertussis* (wP) was substituted in many countries by an acellular subunit vaccine (aP) to reduce side effects. Recent epidemiological studies have shown that aP vaccination in infancy induces less durable immunity than wP vaccination. To determine immunological differences associated with aP vs. wP priming, we performed system-level profiling of the immune response in adults primed with aP vs. wP vaccine in infancy following the Tdap booster vaccination as a surrogate to antigen encounter *in vivo*. Shared immune responses across cohorts were identified, including an increase of the blood monocyte frequency on day 1, and strong antigen-specific IgG response seven days after boost. Comparing aP and wP primed individuals, we found a subset of aP-primed individuals with higher levels of expression for several genes including CCL3 on day 3 and NFKBIA and ICAM1 on day 7 post immunization. These observations were supported by increased CCL3 concentrations in plasma of aP primed individuals. Contrary to the wP individuals, the CCL3-high aP subset presented boosted PT-specific IgE responses. Furthermore, higher antigen specific IgG4 and IgG3 antibodies against specific vaccine antigens at baseline and post boost of aP individuals was observed, suggesting a long term maintained difference in the IgG subtype response. Overall our findings demonstrate that, while broad immune response patterns to Tdap boost overlap between aP and wP primed individuals, a subset of aP primed individuals present a divergent response. These findings provide candidate targets to study the causes and correlates of waning immunity after aP vaccination.

## Background

A *Bordetella pertussis* vaccine containing whole killed bacteria was introduced in the mid-20^th^ century and since then demonstrated its efficacy to confer immunity against the pathogen. For decades this vaccine was administered in a mix with toxoid antigens derived from the causative agents of diphtheria and tetanus called DTwP. The pertussis compounds in this vaccine termed (“w” for whole-cell, also wP for short) were substituted by acellular vaccines containing up to five individual protein antigens. This vaccine cocktail was widely adopted in many countries in the 1990s-2000s for primary (DTaP) or booster (Tdap) vaccination (“a” for acellular, also aP for short) because of significant reduction in side effects compared to DTwP. Though the DTaP/Tdap vaccine was shown to induce protection in infants (*1, 2*), questions were raised about its ability to induce long lasting protection (*3-5*) and prevent transmission (*6-8*). A number of countries experienced an alarming increase of whooping cough cases (*9-11*), leading to a concern that usage of acellular vaccines might be the cause of recent outbreaks (*12*).

Attempts, including from our own group, were undertaken to characterize and compare immunological response in DTwP and DTaP primed individuals in terms of humoral (*4, 13-15*) and T-cell mediated immunity(*16-20*). These studies showed that both wP and aP-primed individuals are capable of strong humoral response resulting in high levels of IgG against *B. pertussis* antigens. However, T cell phenotypes and polarization differed, and those differences persisted decades after the prime vaccine (*21-23*), which has been suggested to be linked with dissimilar vaccine efficacy and immune-imprinting (*23-26*).

While several studies were undertaken to compare immune response to booster vaccination with Tdap in aP- or wP-primed individuals (*17-19, 27, 28*), each of these studies were focused on specific immune responses in isolation. Here we wanted to apply a diverse set of assays to probe the immune response development post vaccination at the systems level. Such approaches have previously been applied to characterize vaccine responses against influenza (*29*), herpes virus (*30*),malaria (*31*) and others (*32*). We wanted to utilize this systems approach to characterize the response to Tdap boost of differently primed individuals. Vaccinated individuals were profiled on transcriptomic, proteomic, and blood cell composition levels during the first two weeks after booster vaccination, and humoral responses were profiled over a three-month period. This set-up allowed us to uncover shared signatures marking the different stages of immune responses to Tdap boost as well as subsets of individuals that showed divergent response patterns.

## Results

### 1. Subject recruitment and study design

To study the long-term effect of priming with the aP vs. wP vaccine, we recruited individuals born in the US prior to 1995, which will have been primed with the wP vaccine in infancy, vs. individuals born later, which will have been primed with aP. The recruited individuals were eligible for booster vaccinations with Tdap, containing tetanus toxoid (T) diphtheria toxoid (d) and acellular Pertussis (ap; FHA, Fim2/3, PRN, and PtTox) antigens. We collected longitudinal blood samples prior to booster vaccination (day 0) and at days 1, 3, 7 and 14 post vaccination. A subset of individuals gave samples 1 and 3 months post vaccination. These samples allowed studying how the immune system responds to pertussis antigenic challenge *in vivo* as a proxy of infectious challenge, and whether this response differed in aP vs. wP primed individuals 15 years or more after the original vaccination. To provide a system level view of the vaccine induced immune response, we set out to identify perturbations at the level of 1) gene expression in whole PBMC 2) cell subset composition and activation states by mass cytometry (CyTOF) 3) protein content in plasma and 4) vaccine specific antibody titers and isotypes in plasma. A total cohort of N=58 donors was recruited, and different assays were performed on subsets of these donors as summarized in **Figure 1** and broken down per individual in **Table S1**.

**Figure 1.**
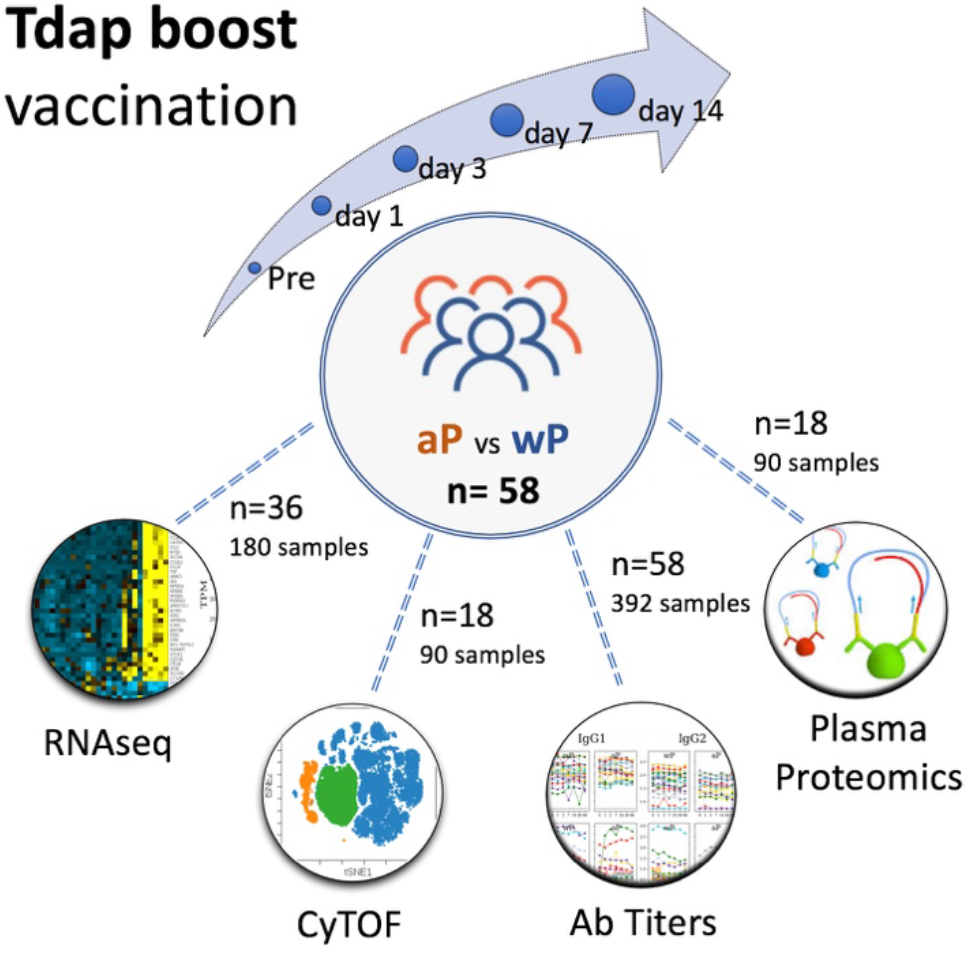
Outline of recruitment and study design. A total of N=58 subjects were enrolled, and blood samples were collected prevaccination (day 0), and at day 1, 3, 7 and 14 following booster vaccination. In addition, plasma was collected at 1 and 3 months postvaccination. Four different sets of assays were performed: Gene expression by RNA-Seq in PBMC, protein marker expression by CyTOF, vaccine specific antibody titers by Luminex assay and plasma protein concentration by PEA (proximity extension assay).

### 2. Tdap boost induces perturbations in PBMC gene expression that follow distinct kinetic patterns

We performed RNA-Seq analysis on PBMC samples collected longitudinally at baseline and following booster vaccination. After removing samples not passing quality controls, complete time courses were obtained for N=36 individuals (16 aP vs 20 wP). To reduce the dimensionality of the obtained gene expression data, we first performed an unbiased clustering analysis that grouped genes together that were co-expressed across samples from different time points and different donors (see methods for more details). This identified 39 clusters of genes (annotated as TrC1…TrC39, **Tr**anscriptomic **C**lusters, **Table S2**). For each sample, the expression level of genes in each cluster was quantified using a principal component analysis, where the first principle component (PC1) was taken as a proxy for expression of genes in the module. Inspecting the expression level of clusters at different time points after booster vaccination revealed that some clusters had essentially unchanged expression over time, while others showed clear perturbations (**Figure S1**). To quantify which clusters were significantly perturbed by the booster vaccination, we picked the time-points of highest and lowest expression for each cluster, and then determined if the difference in expression between these two points was statistically significant using the paired, non-parametric Wilcoxon test (**Table S3**). We separately tested clusters for all individuals, only aP-primed individuals, and only wP-primed individuals. This identified that 28 of the 39 gene clusters showed significant perturbations after vaccination in at least one of the comparisons made.

The 28 perturbed gene clusters were grouped together based on similar expression kinetics using agglomerative clustering (**Figure S2**) into five groups of clusters whose expression kinetics are shown in **Figure 2**. The first group consisted of ten clusters (2346 genes in total) and was characterized by an early increase after vaccination at day 1 and 3. Group 2 consisted of two clusters (246 genes in total) with a distinctive early peak of expression at day 1. Group 3 consisted of three clusters (262 genes in total) that had a significant drop in expression at day 1, which recovered nearly to baseline at day 3. Group 4 included 12 clusters with a substantial peak at day 7 which comprised 2360 genes in total. Group 5 consisted of a single cluster that was significantly perturbed only in wP-primed subjects, and showed increases in expression starting at day 3. Overall this showed that Tdap booster vaccination induced significant perturbations in gene expression of PBMC as detected by RNA-Seq, and that these perturbations followed distinct kinetic patterns.

**Figure 2.**
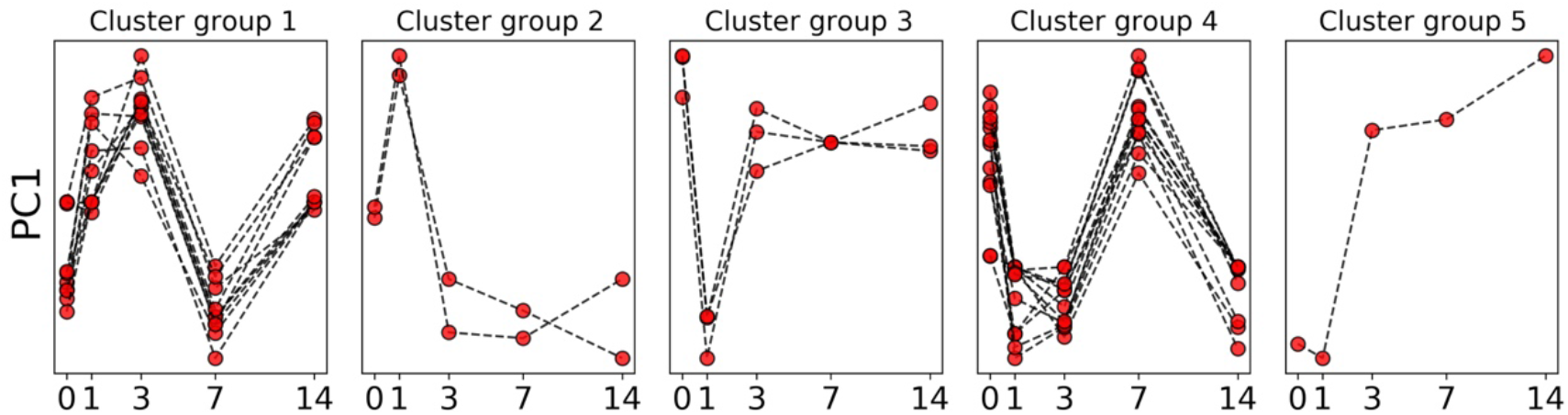
Kinetic groups of transcriptomic clusters. Every group consists of several transcriptomic clusters. Every line indicates min-max normalized PC1 of an individual cluster.

### 3. Tdap boost induces changes in PBMC cell composition that correlate with changes in gene expression

PBMC are a heterogeneous mixture of different cell types that vary substantially in frequency between individuals even in the absence of immune perturbations (*33*). Differences in cell type frequencies in PBMC across individuals are expected to impact gene expression patterns. We systematically analyzed variations in cell type frequencies across different time points for N=18 donors (8 aP vs 10 wP-primed donors) using a CyTOF panel (**Figure S3**). High-dimensional automated gating analysis using the DAFi method (*34*) allowed identification of 21 predefined/distinct cell types and their frequencies, including cell populations that are difficult to resolve by manual gating analysis such as memory T cells and Tregs (**Figure S4**). We determined which of these cell types showed significant perturbations post boost in all individuals, or for either aP- or wP-primed individuals separately. This identified a total of 12 cell populations that showed perturbations (**Figure 3**), including myeloid cells (classical and intermediate monocytes, mDCs) which increased at days 1-3 post boost, and antibody-secreting cells (ASCs) which peaked at day 7. NK cells were characterized by a steady increase through the 14 days of observation. Most of the T cells subsets with significant perturbations showed a drop at day 1, day 3, or both (CD4+ Tcm, CD4+ Tem, CD8+ Tem, CD3+ T cells, CD8+ T cells), while two T cell populations (CD4+ Temra and Tregs) instead showed a slight increase at day 1 and/or day 3. Overall, similar to the gene expression data, we found a heterogeneous set of kinetics upon Tdap boost associated with different cell types.

**Figure 3.**
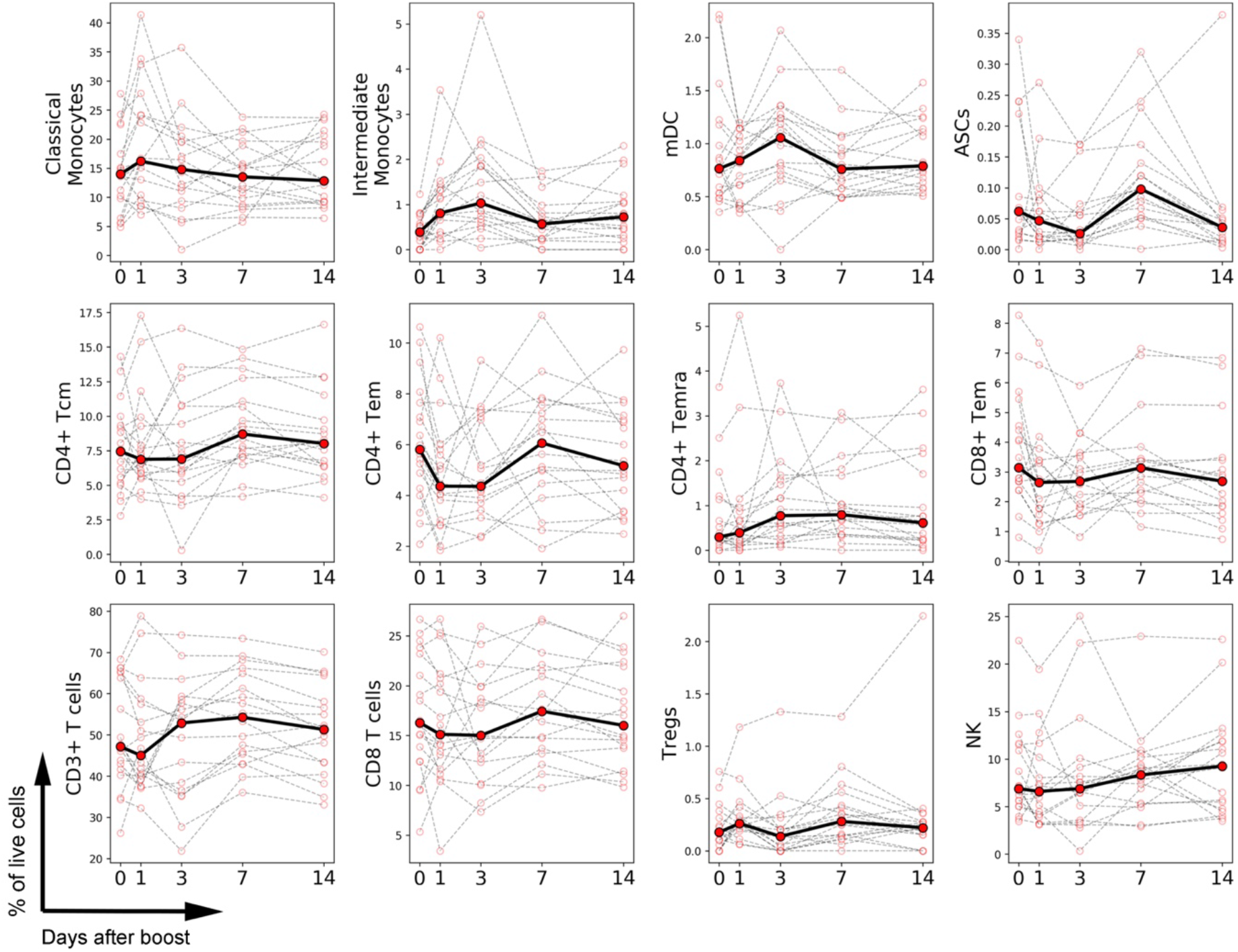
Frequency of cell types altered by vaccination as determined by CyTOF.

Each plot represents the frequency of a given cell type in terms of percentage of live PBMCs (y axis) as a function of the time post booster vaccination (x-axis) as determined by high-dimensional automated gated analysis of data generated by CyTOF. Data is expressed as connected time points from day 0 to day 1, 3, 7 and 14 post for each individual donor (thin lines) and the median of all (bold line). Only cell types that showed a significant difference in frequencies over time are shown, based on a paired, non-parametric Wilcoxon test.

Next, we correlated the frequency of cell types determined by CyTOF with the expression level of gene clusters identified by RNA-Seq for the N=90 samples (18 subjects × 5 time points) where both RNA-seq and CyTOF data was available (**Table S4**). We found several gene clusters whose expression was highly correlated with cell type frequencies (**Figure 4**). This included cluster TrC24 which correlated (Spearman r = 0.71) with frequency of B cells across samples. TrC24 contains the B cell lineage marker CD19, further suggesting that its expression is directly linked to the frequency of B cells in each sample. Notably, the gene cluster shows no significant perturbation in average expression over time post boost and neither does the B cell frequency. Thus, the different expression levels of genes in the TrC24 cluster reflect different baseline frequencies of B cells across individuals, ranging from 5 to 20% of PBMCs, which were not systematically impacted by the booster vaccine. In contrast to B cells overall, cluster TrC27 was significantly correlated with the frequency of antibody secreting (B) cells (ASCs, Spearman r = 0.58). Of the 49 genes in TrC27, 38 code for immunoglobulin heavy and light chains. Both expression of this cluster and frequency of ASCs peak at day 7, which is in line with previous reports of a peak in antibody production at day 7 post booster vaccination in other systems such as influenza (*29, 35*). Thus, Tdap booster vaccination does not perturb the overall level of B cells in PBMC, but it does increase the frequency of antibody secreting cells which peak at day 7, corresponding with the kinetics of the group 4 gene cluster.

**Figure 4.**
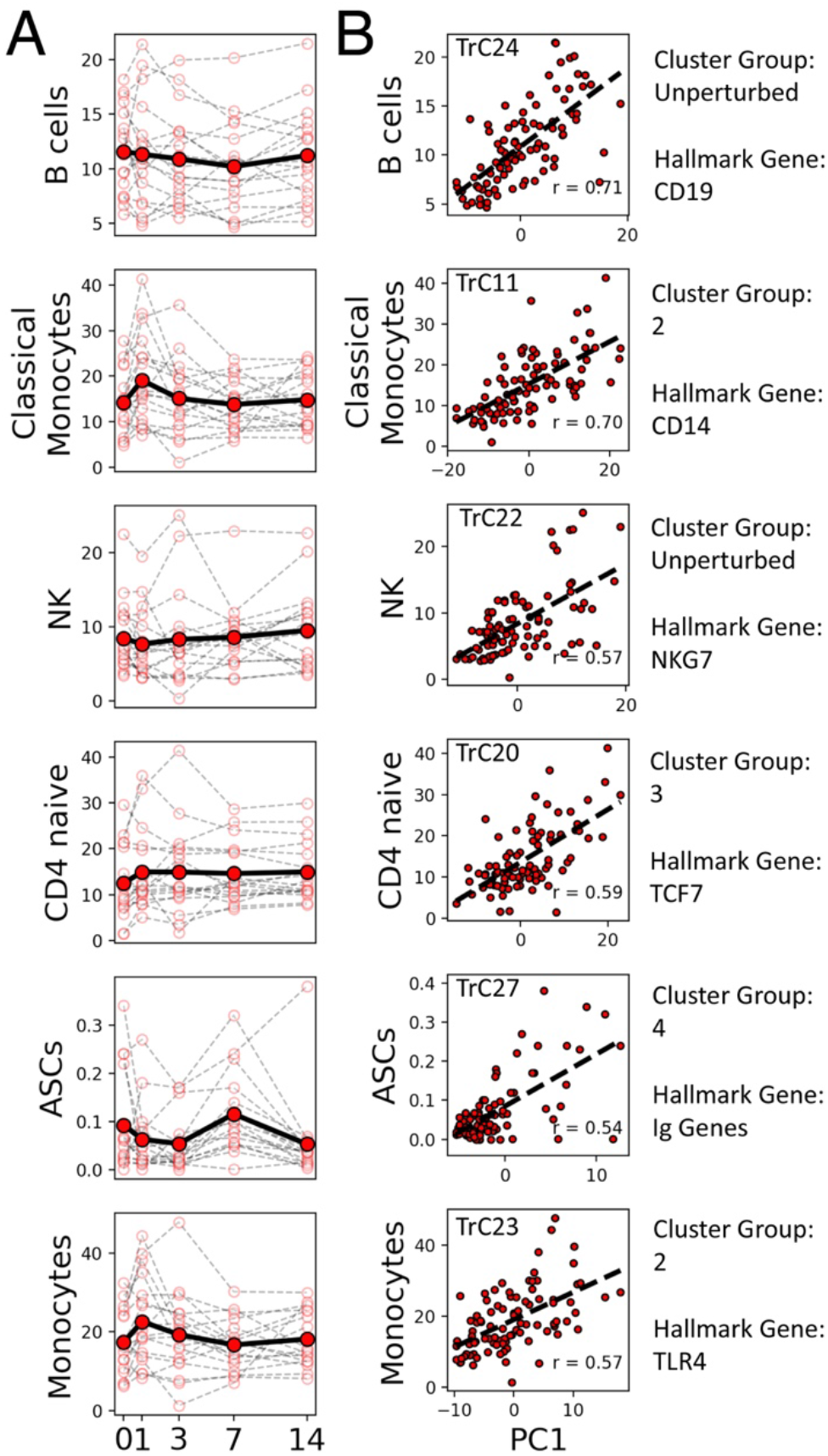
Frequencies of several cell types correlate highly with gene expression levels of specific clusters. The plots show all cell types for which correlations of r>0.5 were found with a specific gene cluster, namely B cells, classical monocytes, NK cells, CD4 naïve cells, ASCs and monocytes. **A**) Frequency of the cell type % of live cells are plotted as a function of time post booster vaccination. Open circles and thin lines connecting them indicate individual responses. Closed circles and the solid lines connecting them indicate average responses over all donors. **B**) Correlation of cell type frequencies with gene expression of the best-matching RNA-seq cluster (different cluster for every cell type), quantified by the first principal component (PC1). The specific cluster is indicated on the top left corner of each plot, its cluster group, and a hallmark gene are indicated on the right.

The second highest correlation for gene cluster expression and cell type frequency was observed for classical monocytes and the gene cluster TrC11 (r = 0.70, **Figure 4**). The TrC11 cluster contains CD14, the lineage marker of classical monocytes, and falls into cluster group 2, which peaked in expression at day 1 post vaccination. This cluster also contains the LYZ- and the S100A9 gene, which distinguish classical from non-classical monocytes. The parent population of total monocytes also correlated significantly with the group 2 transcriptomic cluster TrC23, which contains genes shared in both classical and non-classical monocytes such as TLR4. Overall, these data suggest that the peak of expression of multiple genes at day 1 post booster vaccination resulted from an increase in the frequency of monocytes in PBMCs, of which classical monocytes make up the vast majority. The next two cell types that correlated highly with specific gene expression clusters were 1) NK cells, which showed high correlation with TrC22. While the perturbation in NK cell frequency was significant, the perturbation in expression of TrC22 was below the threshold of significance, which resulted in it being classified as ‘unperturbed’. The gene cluster did contain NKG7 though, which is an NK signature gene, so the correlation is still likely to be biologically meaningful. 2) Naïve CD4 T cell frequency correlated strongly with TrC20. While the perturbation of the cell frequency was not significant, the perturbation of the gene expression cluster did reach significance. The cluster contained TCF7, a hallmark gene of CD4 T cells. As for NK cells above, this suggests that the correlation between these cell frequencies and the gene cluster are likely to be biologically meaningful.

### 4. Tdap boost elevates selected cytokine concentrations in plasma

To determine how Tdap vaccination impacts proteins secreted into blood, we employed a proteomics approach to uncover vaccination related perturbations of 241 unique plasma protein markers (listed in **Table S5**). N=90 plasma samples (18 subjects × 5 time points) were profiled (8 aP vs 10 wP-primed donors) and evaluated for expression of proteins using a highly sensitive and quantitative proximity extension assay (PEA) (*36*). In order to study the kinetics of cytokines and other factors in plasma, we performed a similar dimensionality reduction approach as we did for gene RNA expression (**Figure S5**) and obtained 8 protein clusters of 5 or more proteins (**Figure S6**) with 209 proteins that presented perturbation across the study (**Table S6**). We then examined if any of these clusters were significantly perturbed over time in all individuals or only in aP- or wP-primed individuals. We found that only one cluster (PrC07) was significantly perturbed, which contained proteins encoded by MASP1, TNFSF14, CCL3, CCL4, HGF, CLEC6A, CXCL8, CLEC4D and CLEC4C. This cluster showed significant increased protein levels at days 1 and 3 after booster vaccination and a return to baseline levels afterwards. When comparing the expression of the PrC07 cluster to gene transcriptomics and cell type frequencies, we found that PrC07 positively correlated with 3 transcriptomic clusters (TrC09, TrC10 and TrC31) and negatively correlated with 7 clusters (TrC03, TrC5, TrC7, TrC13, TrC28, TrC34, and TrC39) all with an absolute Spearman R value of >0.5. All the transcriptomic clusters positively correlated belonged to cluster group 1 (peaking at days 1 and 3) and 6 out of 7 clusters it negatively correlated with belong to cluster group 4 (upregulated at day 7). Overall, this showed that some protein concentrations in plasma had a distinct and transitory kinetic response to booster vaccination.

### 5. Tdap boost induces an isotype- and antigen specific pattern of antibody production with transcriptomic changes preceding increased antibody titers

To characterize the humoral response to Tdap, we first profiled the expression of immunoglobulin genes in PBMC transcriptomic data. RNA-seq reads were mapped to the constant regions encoded by IGHG1-4, IGHE, IGHM and IGHD genes, followed by TPM transformation. As shown in **Figure 5A**, RNA expression of all IgG heavy chains peaked at day 7, and a similar pattern was observed for IgM. Transcripts of the IgD heavy chain, on the contrary, did not demonstrate a profound increase at day 7, and the increase in IgE transcripts showed a smaller, but not significant increase.

**Figure 5.**
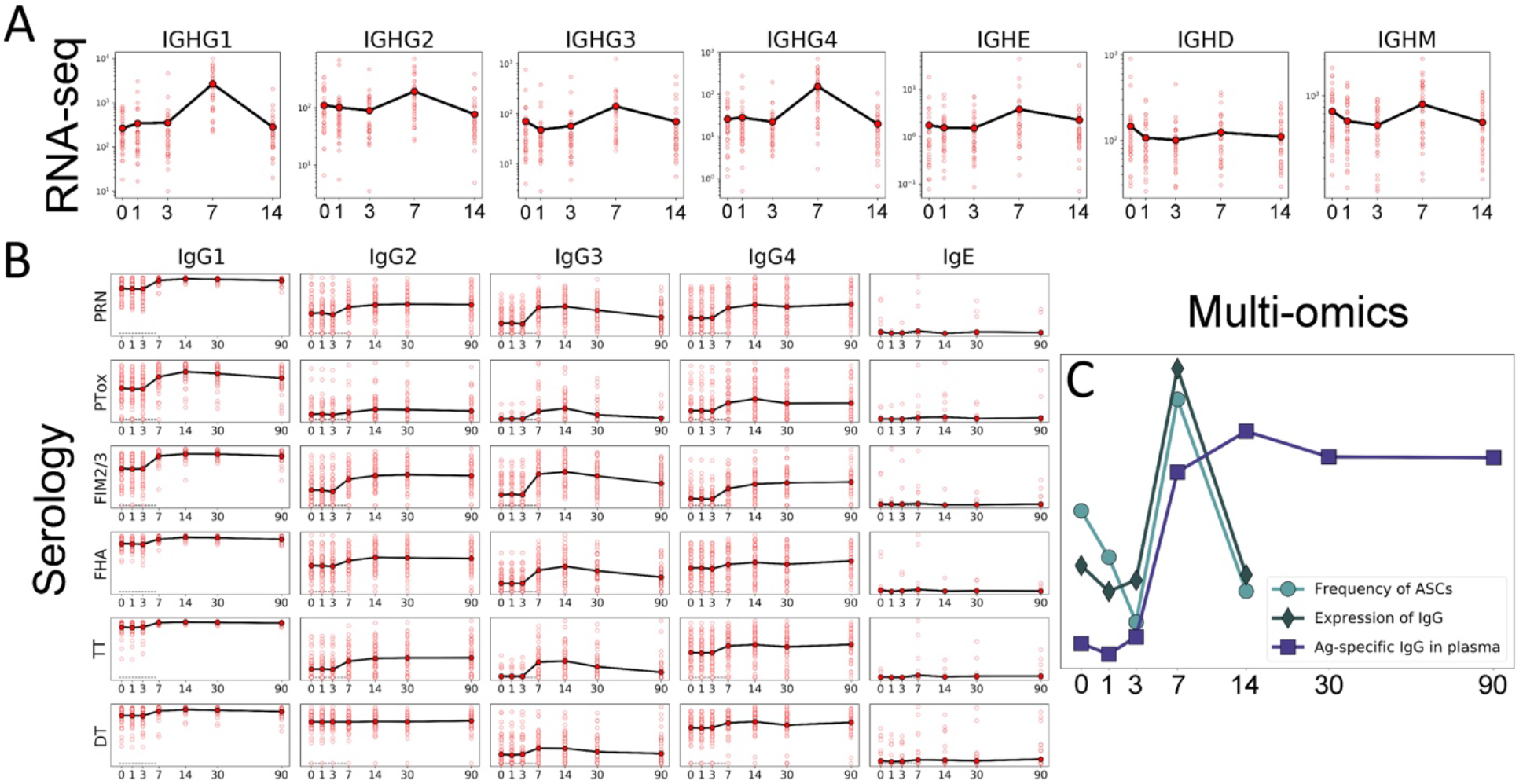
Longitudinal patterns of humoral response. In all plots X-axis denotes days after vaccination. Averaging is done within all cohort. **A**) Expression of immunoglobulin constant regions from RNA-seq **B**) Tdap-specific IgG subclass titers (log10-scaled) from plasma. **C**) Overlay of values from RNAseq, Ab and CyTOF combined measurements. Average of PC1 across all the profiled donors is plotted for RNA-seq and IgG titers data as well as the cell frequency specific for ASCs.

We next profiled the plasma levels of antigen-specific IgG, all IgG subclasses (IgG1-4) and IgE in the same kinetic fashion as before (1 to 14 days) and up to 3 months after boost. Immunoglobulins specific for the antigens contained in the Tdap vaccine were assessed, namely PT, PRN, FHA and FIM2/3. As a control, two whole-cell vaccine antigens not contained in Tdap (adenylate cyclase and lipopolysaccharide of *B. pertussis*) were also profiled, and as expected, no increase of wP specific-IgG levels was observed post booster vaccination (**Figure S7**). In contrast, we observed strong increases in all IgG subclass levels against antigens contained in Tdap (**Figure 5B**) starting at day 7 of the observation period. The peak of antigen-specific IgG levels was observed on day 14 whereas by day 30, the immunoglobulin levels started to decrease. Interestingly, IgG3 levels decreased the fastest (**Figure 5B**). Tdap-specific IgG antibodies were induced in an-antigen- and IgG subclass-specific fashion. For example, there was almost no IgG2 response against diphtheria toxoid (**Figure 5B**). The strongest responses were observed against FIM2/3 antigen (especially IgG1 and IgG3). Overall when averaging ranks across antigens, the IgG1 response was the strongest while IgG2 was the weakest. In concordance with immunoglobulin RNA levels, there was no consistent induction of IgE expression.

An overlay of three signals related to antibody production is depicted in **Figure 5C**, namely the detection of antigen specific IgG in plasma, the RNA expression levels of IgG, IgE, IgD and IgM genes in PBMC and the frequency of antibody secreting cells in PBMC by CyTOF. This figure indicates that production of immunoglobulin peaks around day 7 as assessed by gene expression and frequency of antibody secreting cells. The level of antigen specific antibodies in plasma peaks at day 14 and stays elevated until day 90, which is consistent with antibodies having a half-life of several weeks (*37*). Overall, the data provides consistent evidence for an increased influx of antibody producing cells into PBMC peaking at day 7 post booster vaccination, which results in a persistent increase in B. pertussis specific circulating antibodies.

### 6. Differences in the cellular immune response to Tdap boost in wP vs. aP-primed individuals

After characterizing common patterns of immune responses following Tdap booster vaccination, we wanted to identify differences in the response of aP- vs. wP-primed individuals. First, we applied Mann-Whitney U-test to determine if the expression level of any transcriptomics gene module differed significantly between aP and wP individuals at different time points (**Table S7**). Three clusters showed significantly different expression at specific time points: TrC27, TrC31 and TrC38. As stated above, the TrC27 cluster correlated strongly with the frequency of antibody secreting cells. The cluster showed higher expression in aP vs wP individuals at day 0, prior to booster vaccination, so these differences cannot be clearly linked to the Tdap boost response, and were therefore not examined further.

Clusters TrC31 and TrC38 showed differences in expression between aP and wP primed individuals at day 7 post boost (**Figure 6A**). To determine what specific genes in these clusters contribute to the difference in aP vs. wP primed individuals at day 7, we performed differential expression analysis using DESEQ2, limited to the genes in these two clusters. This identified a total of 14 genes with p_adj_<0.05, four from TrC31 the most pronounced of which was ICAM1, and ten from TrC38 with the most pronounced being the NFKB inhibitor protein NFKBIA. **Figure 6B** shows a heatmap of the expression of these 14 genes in the overall cohort at day 7. This highlights that the differences between aP and wP individuals were due to a subset of aP individuals that showed high expression of most of the 14 genes compared to any of the wP donors.

**Figure 6.**
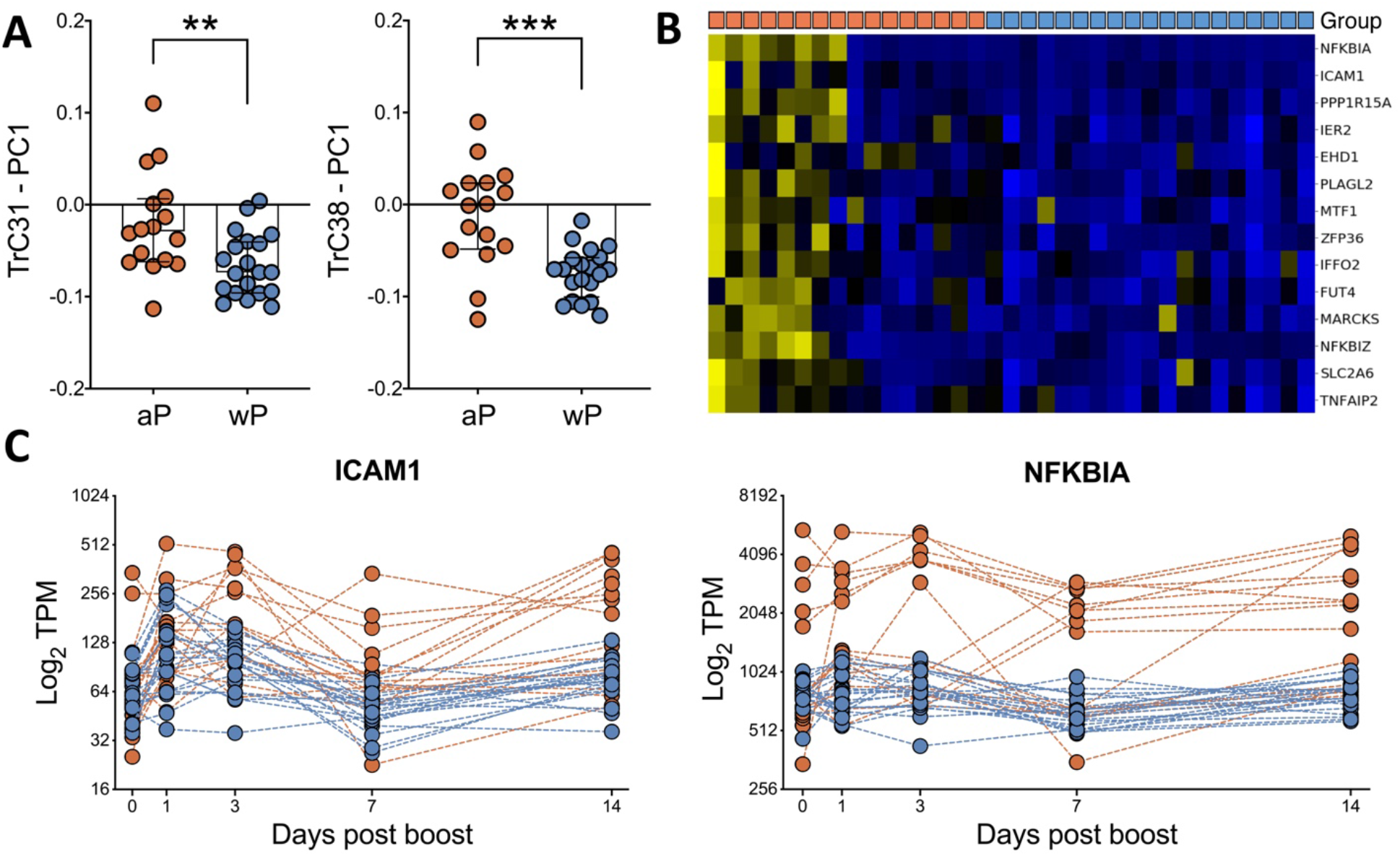
Gene expression differences following in aP vs. wP individuals – focus day 7. **A**) Expression of the genes in clusters TrC31 and TrC38 is significantly different 7 days post boost in aP vs. wP individuals as quantified by the principal component analysis (PC1). **B**) Heatmap of 14 genes from clusters Trc31 and 38 that were identified as differentially expressed on day 7 (Benjamini-Hochberg p_adj_<0.05). Columns denote group of individuals originally primed with either aP or wP as indicated by orange vs. blue boxes in the top row **C**) Expression level of ICAM1 and NKBIA over time, as representative genes in clusters TrC31 and TrC38, respectively, that are differentially expressed on day 7 between aP and wP individuals.

Examining the kinetics of ICAM1 and NFKBIA expression as representatives of differentially expressed genes in cluster TrC31 and 38 (**Figure 6C**) revealed that the subset of aP primed individuals that showed enhanced expression of these genes at day 7 also had high expression at other time points. Specifically, three of the four individuals with the highest ICAM1 expression at day 7 had even higher ICAM1 expression at day 3, and six of the nine aP individuals with higher NFKBIA expression than any wP individual on day 7 already had higher expression than wP individuals on day 3. To examine this in more detail, we performed differential gene expression analysis between aP and wP primed individuals at day 3 post boost. A total of 36 genes were identified as differentially expressed with p_adj_ <0.05 between these cohorts, independent of any specific module considered (**Figure 7A**). As expected, aP-primed individuals again had significantly higher expression of NFKBIA and ICAM1 at day 3. In addition, inflammatory cytokines such as CCL3, CCL4, IL6 and TNF were also expressed highly in the same subset of aP individuals.

**Figure 7.**
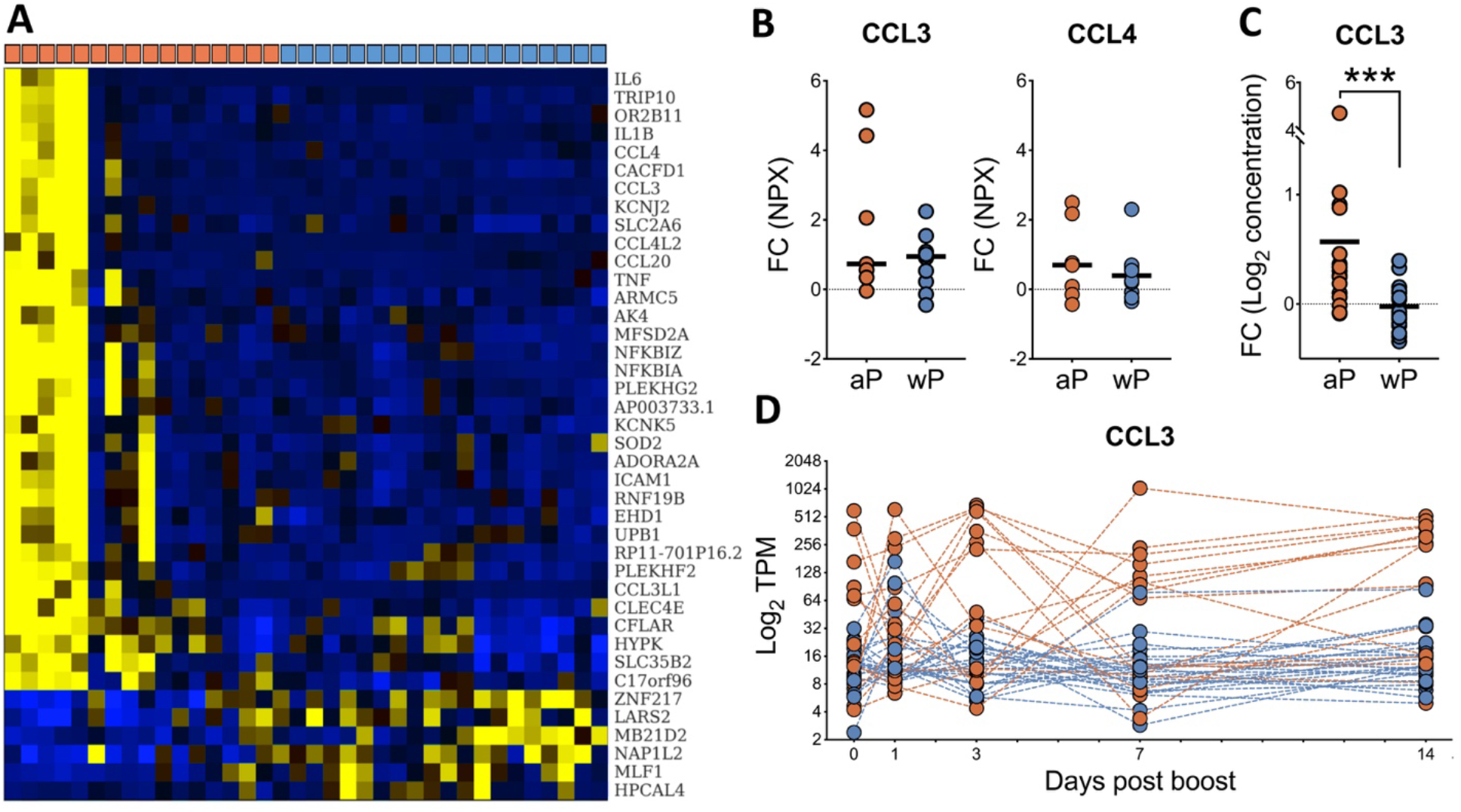
Observed differences in aP and wP-primed individuals – focus day 3. **A**) Heatmap of genes differentially expressed between the cohorts on day 3. Columns denote different individuals primed with aP vs. wP as indicated by orange or blue boxes, respectively in the top row. **B**) Differences in plasma concentration of CCL3 and CCL4 between day 0 and day 3 based on change of NPX values as obtained by PEA assay in human plasma. **C**) Log fold-change from pre-boost to day 3 of plasma concentrations of CCL3 by ELISA. Statistical differences were evaluated using a two-sided, non-parametric Mann-Whitney test (***p=0.0002). **D**) Modulation of CCL3 gene expression over time. aP and wP donors are represented in orange and blue respectively.

To examine if the increased transcription of the cytokines genes corresponded to increased protein expression, we examined the concentration levels of CCL3 and CCL4 in plasma as resulted from proximity extension assay. Indeed, we found increased expression in aP compared to wP individuals (Mann-Whitney U-test, p = 0.012) and this trend was also obtained for CCL4 (**Figure 7B**). To increase the number of individuals for which we had data available, we analyzed CCL3 concentrations using ELISA in plasma samples not evaluated by proximity extension assay (38 donors = 17 wP vs 21 aP), which confirmed that the aP-primed individuals had significantly (p < 0.0001, Mann-Whitney U-test) higher induction of CCL3 in plasma on day 3 than wP-primed individuals (**Figure 7C**). This confirmed through multiple independent assays that a subpopulation of aP individuals showed a distinct cellular immune response profile with enhanced inflammatory cytokine production on day 3.

### 7. Differences in the humoral immune response to Tdap boost in wP- vs. aP-primed individuals

As mentioned above, Tdap specific antibodies were induced by day 7 after booster vaccination. In comparing the total IgG titers against several Tdap antigens between aP and wP cohorts (see **Figure S9**), no significant differences were found. Next, we looked into antigen-specific response at the level of IgG subclasses. After FDR correction, we found that at the peak of the antibody boost on day 14, aP-primed individuals showed higher IgG4 response against FHA and higher IgG3 response against FIM2/3 than wP-primed individuals. Importantly, these higher titers in aP individuals were already observed prior to the booster vaccine, suggesting long-lived differences in the level of FHA specific antibodies of these isotypes between the cohort (**Figure 8A**). The pre-booster titers for the IgG4 subclass antibodies in particular were substantial, so we examined if the difference in IgG4 antibodies prior to boost correlated with any of our measures of the cellular immune response at later time points. The highest correlation we obtained was r = −0.49 for cluster TrC21, which was the sole cluster falling into the kinetic group 5 that peaked on day 14 and was only found significantly perturbed in wP-primed individuals. This cluster contained genes such as the chemokine CXCL5 and several integrins associated with platelets.

**Figure 8.**
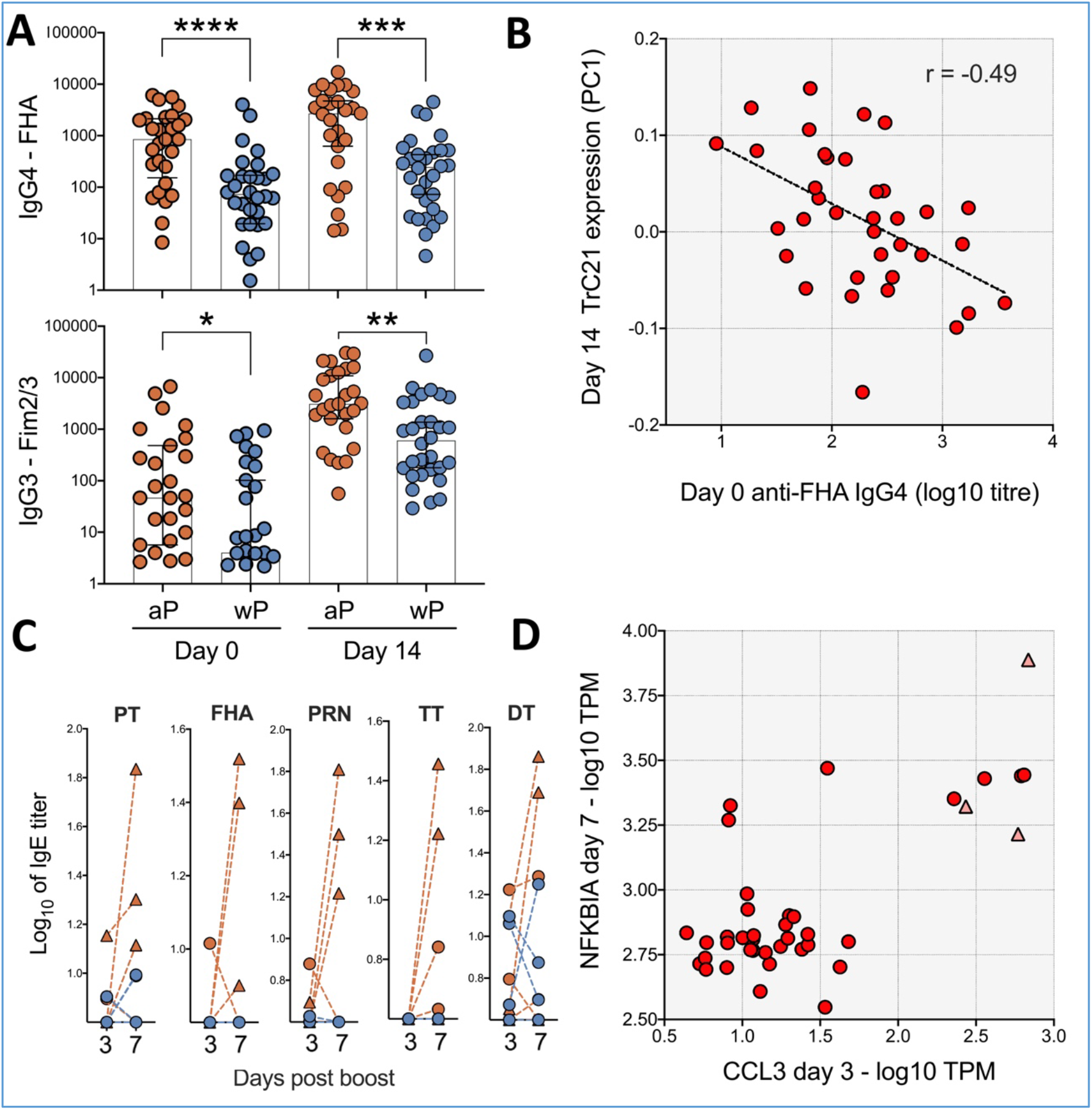
Differences in humoral responses against the pertussis vaccine antigens in aP vs. wP donors. **A**) FHA (top) and Fim2/3 (bottom) prior to booster vaccination (left) and 14 days after (right). Statistical differences were evaluated using a two-sided, non-parametric Mann-Whitney test (****p<0.0001, ***p=0.0002, **p=0.0015, *p=0.0141). **B**) IgG4 antibody titers prior to boosting (x-axis) negatively correlate with the expression of module TrC21 14 days post boost (r = −0.49) **C**) IgE level modulation over time for the several antigens in individuals vaccinated with aP (orange) or wP (blue lines) in infancy. Three individuals from the aP group (triangles) showed consistent increases of IgE from day 3 to day 7 post booster vaccination. **D**) Scatter plot of RNA-Seq gene expression data of NFKBIA on day 7 (y-axis) v. CCL3 expression on day 3 (x-axis). The three individuals with consistent IgE responses are marked with red triangles.

We further analyzed antigen-specific IgE responses, which have previously been reported as being enhanced in aP individuals (*14, 38, 39*) especially with underlying atopic conditions (*40*). The majority of subjects did not demonstrate any IgE increase at day 7, but three out of thirty aP-primed individuals (10%) did possess a significant increase in IgE (at least by 50%) levels against 3 or more Tdap antigens (**Figure 8C**). For wP-primed individuals, IgE responses were never consistent across antigens: two donors responded with IgE against diphtheria toxoid only and one donor against tetanus toxoid (non-pertussis antigens). As a control for allergic status of the donors, we also looked at IgE response against 2 seasonal allergens. We observed two potentially allergic donors in aP cohort and 2 potentially allergic donors in wP cohort, which suggested that the tendency for a higher IgE response in aP individuals was not driven by an increased presence of allergic individuals. Notably, while the number of individuals with an increased IgE response was small, all three of these individuals were also among the high CCL3 expressers on day 3 and high NFKBIA expressers on day 7 (**Figure 8D**). This suggested that the differential gene and protein expression patterns observed in a subset of aP individuals might be linked with differential antibody isotype polarization.

## Discussion

Whooping cough is a highly contagious disease caused by *Bordetella pertussis*. Since the introduction of the vaccine against its agent in the mid-20^th^ century infection rates dropped dramatically for the US reaching as low as <1 per 100’000 in the 70s (*41*). For decades inactivated *B. pertussis* was supplemented by diphtheria and tetanus toxoids to formulate DTwP vaccine. Due to concerns about safety of this vaccine containing bacterial cells it started to be replaced with DTaP where *B. pertussis* antigens are constrained to only a few proteins (*42*). In the recent decade outbreaks of whooping cough occurred in several countries and overall infection rates also increased (*43*). Many factors can contribute to this increase such as reduction in vaccination coverage and increased surveillance. However, adoption of the acellular vaccine and the differential immunity elicited is often considered as one of the most significant contributors (*12*). In this study, we aimed to utilize multisystem profiling of immunological response to Tdap boost in wP- and aP-primed individuals to uncover immune signatures of the booster immunization that are shared across individuals, and those that differ based on the type of the priming vaccine.

In terms of shared immune signatures, we found significant alterations in the transcriptomic profile on days 1 and 3 post-boost, reflecting innate and early onset of adaptive immunity. Several of the observed changes in the transcriptomic signatures could be explained by changes in the cell composition in PBMC, which showed a marked increase in the frequency of monocytes on day 1 and 3, and an accompanying decrease of other cell types. Of note, given that RNA-Seq data measures relative concentrations of gene expression in PBMC, and given that the cell frequency data we obtained is a proportion of total live cells, it is possible that most of the changes observed are explained by an absolute increase in the release of monocytes into the bloodstream following vaccination, which has been reported before (*44*). Such an absolute increase of monocytes would also result in a decrease of the relative frequency of other cell types, such as T cells.

Beyond the vaccine induced alterations in transcription on days 1 and 3 that could be explained by cell type composition of PBMC, we also found transcriptomic perturbations that were indicative of cell activation. Several of these changes were accompanied by a steady increase in detection of proteins in plasma including CCL3, TNFSF14, CXCL8 and several C-type lectins. CCL3 is a pro-inflammatory cytokine affecting many immune cells (*45*). It is also known to induce Th1 response and affect T cell differentiation (*46*). Tumor necrosis factor superfamily ligand TNFSF14 (also known as LIGHT) is regulatory molecule which can be produced in membrane-bound as well as soluble form. It can affect inflammatory as well as structural cells (*47*). It can co-stimulate T-cell activation and proliferation (*48*). CXCL8 (IL-8) is another proinflammatory cytokine involved in neutrophil recruitment (*49*). Induction of CXCL8 secretion by PMBCs was observed in study of Shingles vaccine (*30*). CCL3 and TNFSF14 were not reported in that or other studies of antiviral vaccines (*29, 37*)and potentially highlights a specific response to Tdap booster vaccination.

On day 7 post boost, we observed significant perturbations across our readouts that were consistent with a peak of the induction of the adaptive humoral response. We observed a peak in the transcription of antibody heavy chain genes in PBMC based on RNA-Seq coinciding with a peak of the frequency of antibody secreting cells in PBMC as measured by CyTOF. This presumed peak in antibody production was accompanied by a significant increase of vaccine specific antibody titers in plasma, which peaked later (on day 14) as expected due to the longer half-life of circulating antibodies resulting in continued increase of antibody titers as long as antibody secreting cells are active.

In addition to the similarities of the responses to Tdap boost in aP- vs. wP-primed individuals, we also noted a number of differences. In terms of humoral responses, while there were no differences in overall IgG titers, there were differences in an IgG subtype and antigen specific fashion: aP-primed individuals possessed higher IgG4 response against FHA and IgG3 response against FIM2/3. This difference was observed both at the peak of the response on day 14 and on the day prior to vaccination suggesting an imprinted memory B-cell response from prior vaccination. Given that the antibodies of the IgG4 isotype have limited or no opsonization function, its higher levels against the FHA antigen in aP-primed individuals might be linked with reduced ability to prevent colonization in these individuals (*8*). Moreover, we found a negative correlation between high IgG4 titers for FHA prior to boosting and expression of a gene module 14 days post boost that is significantly induced in wP individuals and contains the chemokine CXCL5 and several genes associated with platelets. While it is tempting to speculate that the two are causally linked, additional studies will be needed to examine mechanistic links between the two. Moreover, the fact that the trends for IgG4 responses were not observed uniformly for all pertussis vaccine antigens complicates the interpretation of these data.

Profiling of IgE responses revealed further differences between aP- and wP-primed individuals: While a subset of aP-primed individuals had substantial induced IgE response against several antigens, no such responses were observed for wP-primed individuals. This observation is in line with the previously reported polarization of aP-primed individuals towards Th2 responses (*16*) which are expected to correlate with increased IgE secretion, and which was also observed in aP-primed infants (*14*). Observing differences in antibody isotype polarization decades after the initial priming is interesting and was also observed at the level of T cell responses (*18*).

In addition to differences in the antibody response, we also found disparities in the cellular response between aP- vs. wP primed individuals, which were driven by subsets of aP-primed individuals that had a higher inflammatory signature higher than other individuals, marked by increased expression of gene expression modules that included ICAM1, NFKBIA and CCL3. The elevated expression of CCL3 in a subset of aP-primed individuals on day 3 post boost was confirmed through multiple assays (RNA-Seq, Proximity Extension Assay, ELISA), and these ‘CCL3-high’ responders also included all individuals that showed boosted IgE responses to pertussis vaccine antigens.

Overall, our study is the first to provide a comprehensive picture of immune responses to Tdap booster vaccination for individuals primed with the aP- or wP-vaccine in childhood. The observed shared immune response to Tdap boost was largely in line with those observed in other vaccine studies (*13, 29, 44, 50*). The differences discovered between aP- vs. wP-primed individuals will require further examination, but in our minds raise two main hypotheses that could explain differences in vaccine efficacy: (1) aP-primed individuals show a higher level of IgG4 antibodies prior- and post-boost as a result of initial priming. (2) a subset of aP-primed individuals shows a differential response marked by high expression of indicator genes (CCL3/4, ICAM1, NFKBIA) post boost that is due to a combination of the initial priming with unknown co-factor(s). Additional studies with samples from separate sets of individuals will be necessary to test and refine these hypotheses.

## Materials and methods

### Study subjects

We recruited 58 healthy adults from San Diego, USA (**Table S1**). All participants provided written informed consent for participation and clinical medical history was collected and evaluated. Clinical data for each patient was collected by multiple approaches. Whenever possible vaccination records were collected from study participants or parents/custodian as appropriate. For some donors, the original clinical vaccine record was not available or incomplete, in which cases information was collected by the clinical coordinators through questionnaires, recording dates and numbers of vaccination, including the information that no boost was administered at least in the previous four years prior to this study. All donors were from the San Diego area, and to the best of our knowledge followed the recommended vaccination regimen (which is also necessary for enrollment in the California school system), which entails five DTaP doses for children under 7 years old (three doses at 2, 4 and 6 months and then two doses between 15-18 months and 4-6 years) and a Tdap booster immunization at 11-12 years and then every 10 years. Individuals who had been diagnosed with B. Pertussis infection at any given time in their life were excluded. Other exclusion criteria were pregnancy at the start of the study; present severe disease or medical treatment that might interfere with study results; any vaccination in the last month and/or antibiotic use or fever (>100.4F (38 C)). In all groups, male and female subjects were included equally and originally vaccinated with either DTwP or DTaP in infancy, received a booster vaccination with Tdap and donated blood, before the boost, 1, 3, 7, 14, 30 or 90 days after the boost. Plasma for the same time point samples was collected after blood processing.

### Booster Vaccination

For booster vaccinations participants received a booster vaccine (Adacel^®^, Sanofi Pasteur, Lyon, France) with Tetanus Toxoid, Reduced Diphtheria Toxoid and Acellular Pertussis Vaccine Adsorbed (Tdap). Each dose of Adacel vaccine (0.5mL) contains the following active ingredients: Detoxified Pertussis Toxin (PT), 2.5μg; Filamentous Hemagglutinin (FHA), 5μg; Pertactin (PRN), 3μg; Fimbriae Types 2 and 3 (FIM); 5μg Tetanus Toxoid (TT); 5Lf; Diphtheria Toxoid (DT), 2Lf. Other ingredients include 1,5mg aluminum phosphate (0.33mg of aluminum) as the adjuvant besides residual formaldehyde, glutaraldehyde and phenoxyethanol. Experimental procedures

### PBMC isolation

Peripheral blood mononuclear cells (PBMCs) were isolated from whole blood by density gradient centrifugation according to the manufacturer’s instructions (Ficoll-Paque Plus, Amersham Biosciences, Uppsala, Sweden) as previously described (*51*). Cells were cryopreserved in liquid nitrogen suspended in FBS containing 10% (vol/vol) DMSO. Alternatively, 6×10^6 PBMC from each sample were transferred directly to Quiazol regent (QIAGEN), resuspended and immediately aliquoted and stored at −80°C until RNAseq downstream processing. Heparin plasma was collected from the upper density gradient layer after blood processing, aliquoted and stored at −80°C.

### Multiplexed Luminex Immunoassays

Antigen-specific antibody responses were measured through a modified multiplexed Luminex assay as previously reported (*18, 40*). Pertussis, tetanus and diphtheria proteins (Pertussis Toxin Mutant; PT, inactivated Pertussis Toxin; 1%PFA PT, Pertactin; PRN, Filamentous Hemagglutinin; FHA, Fimbriae2/3; FIM2/3, Adenylate Cyclase Toxin; ACT, Lipooligosaccharide; LOS, Tetanus Toxoid; TT, and Diphtheria Toxoid; DT from List Biological Laboratory, Campbell, CA and Sigma-Aldrich St. Louis, MO), inactivated Rubeola antigen as an internal vaccine control (Edmonston strain from Meridian Life Science, Inc., Memphis, TN) common allergens as an IgE-allergen control (Cat allergen; Fel d1, Birch allergen; Bet v1, Rye grass allergen; Lol p1 [Indoor biotechnologies, Charlottesville, VA]), irrelevant protein Ovalbumin (OVA, Invivogen, San Diego, CA) and PD1 as an internal negative control were coupled to distinct fluorescent-barcoded MagPlex microspheres (Luminex Corporation). To generate 1%PFA PT, native pertussis toxin (List biologicals) was incubated with formaldehyde for a final concentration of 1% v/v for 1 hr at 4°C. The inactivated toxin was then dialyzed using Zeba^™^ spin desalting columns (ThermoFisher) and protein concentration determined via a micro BCA^™^ protein assay (ThermoFisher). Plasma from each individual or WHO B. pertussis human serum reference standard (NIBSC 06/140 Hertfordshire, UK) were mixed with an equimolar mixture of each conjugated microsphere. The microspheres were then washed with a PBS-tween 20 buffer to release non-specific antibodies and bound antibodies were detected via anti-human IgG phycoerythrin (PE; clone JDC-10) anti-human IgG1-PE (clone HP6001), anti-human IgG2-PE (clone HP6025), anti-human IgG3-PE (clone HP6050, all from Southern Biotech, AL, US), anti-human IgG4-PE (clone HP6025, Abcam, Cambridge, UK) or human IgE-PE (clone BE5, Thermo Fisher) to measure isotype or IgG subclass antigen-specific antibodies. Samples were subsequently analyzed on a Luminex FLEXMAP 3D instrument (Luminex Corporation). PT-, PRN- and FHA-specific IgG positive beads were calculated as IU/mL based on the WHO reference serum. Other antigen-specific IgGs that had no reference standard or antigen-specific IgE, IgG1, IgG2, IgG3, and IgG4-positive beads are reported as log10 of the median fluorescent intensity (MFI).

### Proximity extension assay (Olink PEA)

Plasma samples from 8 aP and 10 wP donors were sent for analysis by Analysis Lab at Olink Proteomics (Olink Proteomics, Watertown, USA) for analysis of a total of 276 proteins using three panels of proximity extension assay (PEA). Briefly, plasma was incubated with paired oligonucleotide labelled antibodies which target specific proteins. Once the antibody recognizes the antigen, the proximity of the oligonucleotide tails allows formation of the DNA amplicon which enables amplification by polymerase chain reaction. Detection of protein-specific PCR products by real time PCR allows collection of Ct values that are then transformed to normalized protein expression units (NPX) which allow comparison of protein expression between samples (*52*). The commercially available PEA panels applied; Immuno-Oncology, Immune Response and Metabolism allowed the identification of a total of 276 proteins and soluble factors in plasma. Two of the aP-primed donors were excluded from the study due to high variability in the technique internal control. A total of 23 analytes were excluded based on the number of samples that presented values below the limit of detection (LOD) to avoid introduction of results bias due to a low number of data points (**Table S8**). Finally, 12 duplicate analytes due to panel overlap remaining after LOD exclusion were evaluated for correlation as internal quality control (**Figure S9**) and only one of the analysis performed in duplicate was included in the final analytes list. The final number of analytes included in the study was 241 for a cohort of 16 donors (6 aP-primed and 10 wP-primed individuals). Those 241 analytes were further filtered by exclusion of proteins with a coefficient of variation over 10% (CV>0.1), which reduced the list of analytes included in the study to 209.

### Enzyme-linked immunosorbent assay (ELISA)

Plasma samples from 20 aP and 19 wP donors was measured by enzyme-linked immunosorbent assay (ELISA). CCL3/MIP-1 alpha DuoSet ELISA and Human LIGHT/TNFSF14 Quantikine ELISA Kit (both from R&D systems) were conducted following the manufacturer’s instructions.

### CyTOF cell analysis

PBMCs were thawed and directly stained with the viability marker Cisplatin followed by a surface antibody cocktail incubated for 30 min. Subsequently, and after washes, cells were fixed in PBS with 2% paraformaldehyde overnight at 4°C. The following day, cells were stained with an intracellular antibody cocktail after permeabilization using saponin-based Perm Buffer (eBioscience). Before sample acquisition and additional washes after staining, cellular DNA was labeled with Cell-ID^™^ Intercalator-Ir (Fluidigm). Samples were kept in the pellet form and resuspend in 1:10 of EQ Beads in 1 ml of MiliQ water and then acquired using a Helios mass cytometer (Fluidigm). Antibodies used in CyTOF are listed in **Table S9**. Automated gating analysis of CyTOF data was conducted using DAFi (Directed Automated Filtering and Identification of cell populations) ^(*34*)^. Source code of DAFi is publicly available at GitHub with configuration and usage information (https://github.com/JCVenterInstitute/DAFi-gating). The DAFi configuration files used in the analysis can be found in https://github.com/JCVenterInstitute/DAFi-gating/tree/Pertussis_CyTOF/LJI_Manuscript for result reproducing. Cell frequencies of all the 21 distinct cell populations were obtained by DAFi with the exception of ASC (antigen-secreting cells/plasmablasts), which was independently analyzed and calculated by manual gating analysis using the combination of CD45+Live+CD14-CD3-CD19+CD20-CD38+ antibody markers.

### RNA-sequencing

RNA sequencing (RNA-seq) were performed as described before (*30, 31*). Briefely, RNA was extracted using the Qiagen miRNeasy Mini kit with on-column DNase treatment and 500 ng of total RNA was used as input for library preparation using the Illumina TruSeq Stranded mRNA Library Prep Kit as previously described. The libraries were sequenced on the HiSeq3000.

### RNA-seq bioinformatics data analysis

Raw sequencing reads were aligned to the hg19 reference using TopHat (v 1.4.1, library-type fr-secondstrand-C) (*53*). Gencode v. 19 (obtained from UCSC Genome Browser) were used for analysis. The HTSeq library htseq-count with union mode was used to quantify reads for all annotated genes. Raw counts were TPM-normalized (Transcripts Per Million reads) for subsequent analysis steps.

### Clustering analysis

For clustering RNA-seq data, weighted gene co-expression network analysis (WGCNA) was performed to identify sets of genes that share a similar expression pattern (*54, 55*). WGCNA parameters were set in the following way: minModuleSize = 20, deepSplit = 4 and MEDissThres= 0. The 8000 most variable genes, that were considerably expressed (max (TMP > 5) among all samples) were considered as input for the WGCNA analysis.

For clustering of proteomics data (PrCXX clusters), only proteins with average intra-donor coefficient of variation greater than 0.1 were considered, i.e. those which were substantially altered after the immunization. Then distance matrix across all samples using correlation metric was computed. ‘Average’ linkage method was applied to construct dendrogram. Threshold for cluster assignment was chosen based on similar approach as for transcriptomic data, finding a trade-off between inclusion of proteins into clusters and cluster tightness (**Figure S6**).

### Function characterization of gene sets

For inferring transcriptional patterns of genes belonging to a cluster the first principal component was extracted using principal component analysis reducing the gene/protein dimension. Gene Ontology enrichment was done in GOnet web application (*56*). Differential expression analysis was done using Bioconductor package DESeq2 (*57*). Genes were considered differentially expressed between aP and wP groups when the DESeq2 analysis resulted in a Benjamini-Hochberg p_adj_ < 0.05.

Plotting was done using Python Matplotlib package.

### Study approval

This study was performed with approvals from the Institutional Review Board at La Jolla Institute for Allergy and Immunology (protocols; VD-101). All participants provided written informed consent for participation and clinical medical history was collected and evaluated.

## Acknowledgments

We are grateful to Jolla Institute for Immunology Flow Cytometry and Bioinformatics core facilities for services, as well as the members of the Peters and Sette laboratories for their help and critical reading of this manuscript. The authors would also like to thank all donors that participated in the study and the clinical studies group staff, particularly Shariza Bautista, Krystal Caluza, Brittany Schwan and Gina Levi for all the invaluable help.

## Author contributions

B.P. and A.S. conceived the study, and R.D.A., F.S. and N.K. conducted the experiments. B.P., A.S., R.D.A., M.P. and F.S., wrote the manuscript. M. P. conducted the bioinformatic analysis and statistical analysis with the collaboration of M.B. and J.B. Y.Q., A.M. and R.H.S. developed DAFi automated gating for CyTOF analysis and conducted the analysis and provided feedback on the manuscript. Y.T. helped with the Ab panel design and operation of the CyTOF mass cytometer. M.C. and B. P. performed and coordinated the RNA-seq experiments and provided feedback on the manuscript. C.D.P., L.A.P. and A.G. run the Multiplex assays to evaluate antibody isotype and antigen specificity and provided feedback on the manuscript. B.P. oversaw all the statistical analysis.

## Data and materials availability

The RNA-seq and other relevant data are in the process of being submitted to Gene Expression Omnibus (GEO) and ImmPort, respectively. Accessions code and study number to be determined.

