## Supplementary Figures for "A system-view of *B. pertussis* booster vaccine responses in adults primed with whole-cell vs. acellular vaccine in infancy"

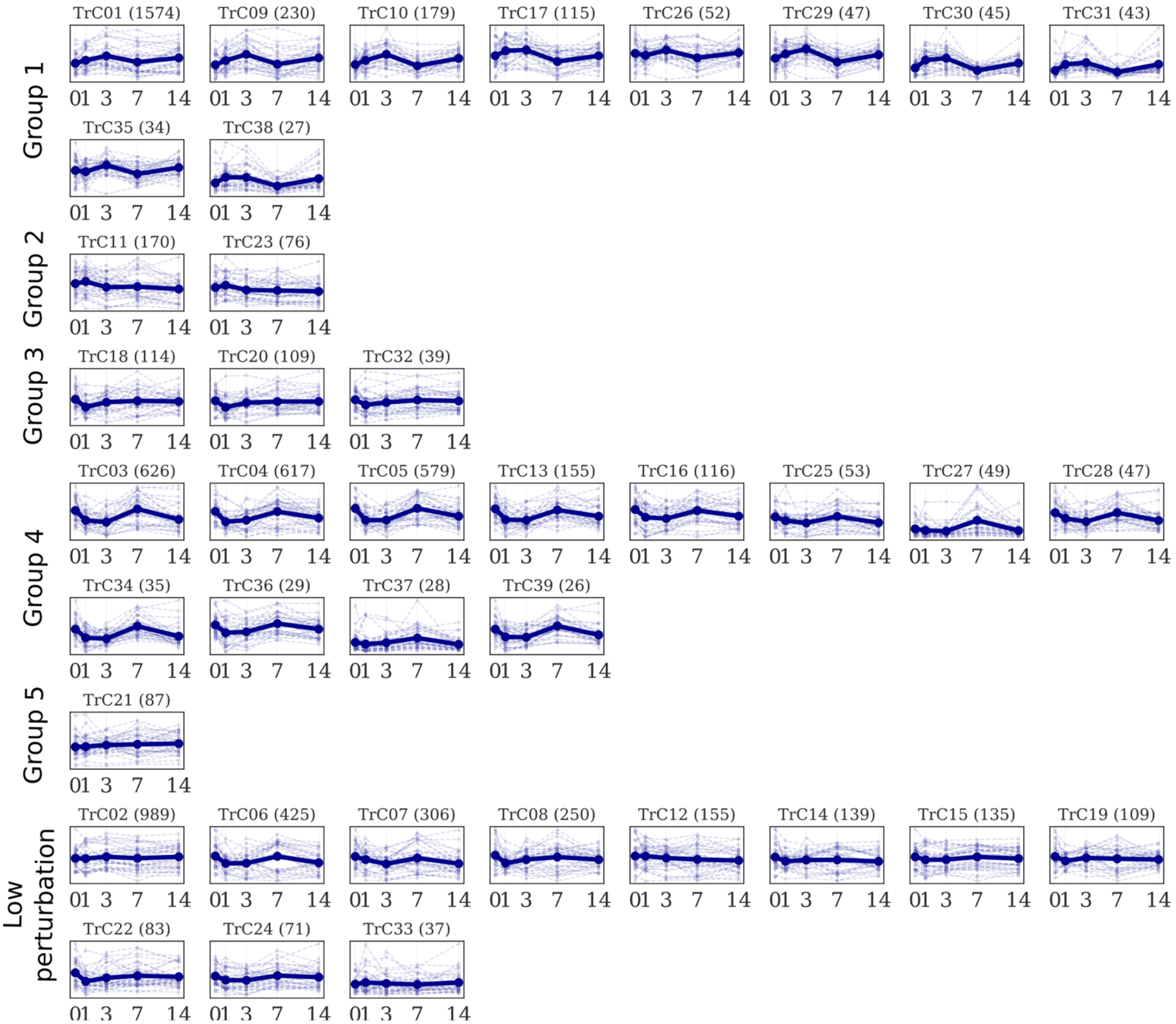


***Fig. S1. Longitudinal patterns of transcriptomic clusters.*** Thick lines represent average of PC1 for a given cluster across all the profiled individuals. Thin lines represent PC1 values for a given individual. Clusters are grouped by longitudinal patterns.

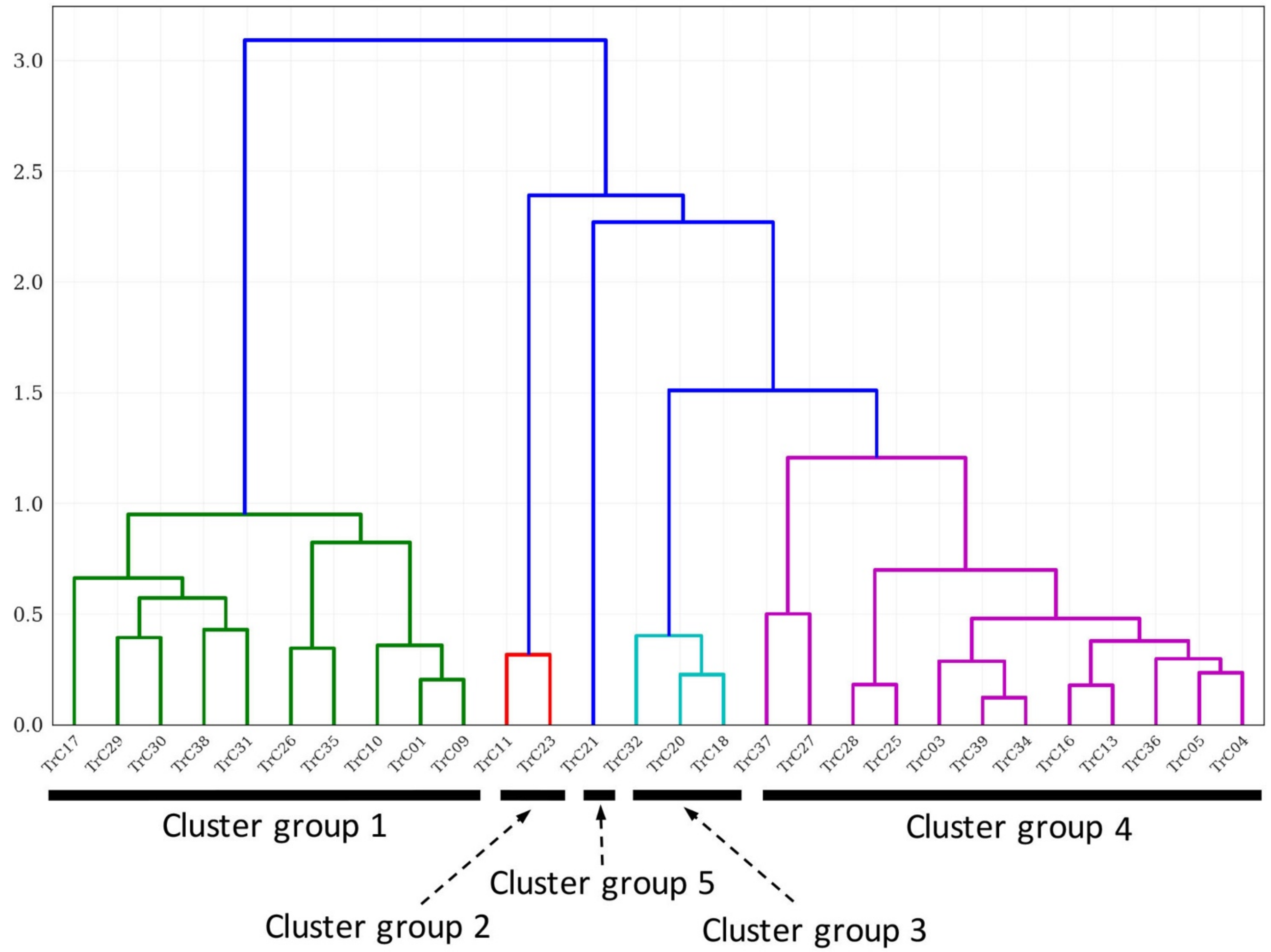


***Fig. S2. Dendrogram for similarity of longitudinal patterns of transcriptomic clusters.*** Dendrogram was constructed using 'cityblock' distance metric and 'average' linkage method.

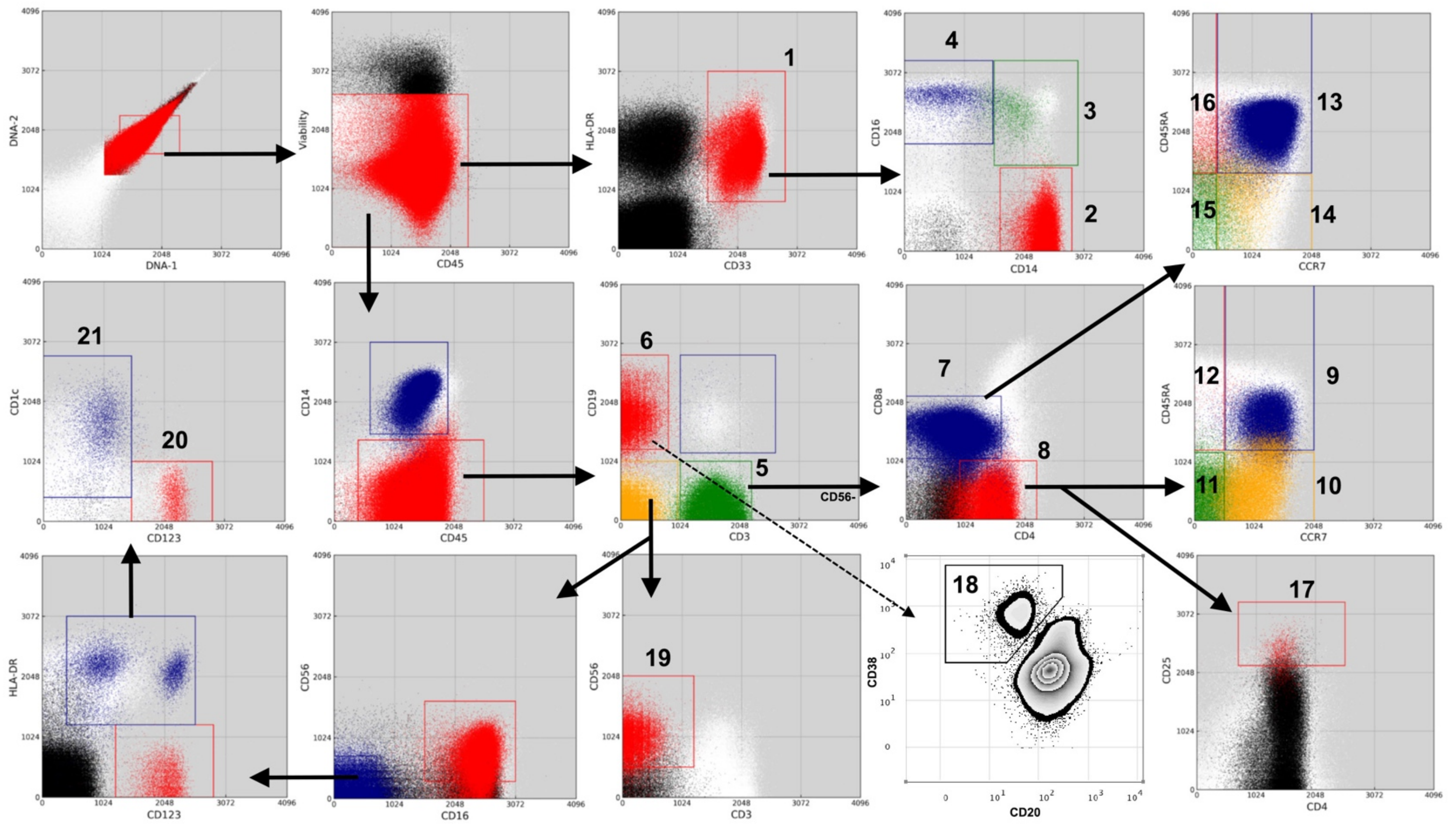


***Fig. S3. Identification of immune cell subsets in human PBMCs by CyTOF.*** Each cell population was identified based on surface protein marker expression following the sequential dot-plot gating strategy by using manual gating analysis (cell population #18) or DAFi automated gating analysis (the other cell populations) . Each number (Gate ID) represents a major population or subset. 1) Monocytes, 2) Classical Monocytes, 3) Intermediate Monocytes, 4) Non-classical Monocytes, 5) CD3+ T cells, 6) B cells, 7) CD8+ T cells, 8) CD4+ T cells, 9) CD4+ naïve, 10) CD4+ Tcm, 11) CD4+ Tem, 12) CD4+ Temra, 13) CD8+ naïve, 14) CD8+ Tcm, 15) CD8+ Tem, 16) CD8+ Temra, 17) Tregs, 18) ASCs (Plasmablasts), 19) NK cells, 20) pDCs, 21) mDCs. Detailed gating strategy and analysis methods are described in Materials and Methods. Representative 2D plots are shown for illustration purpose.

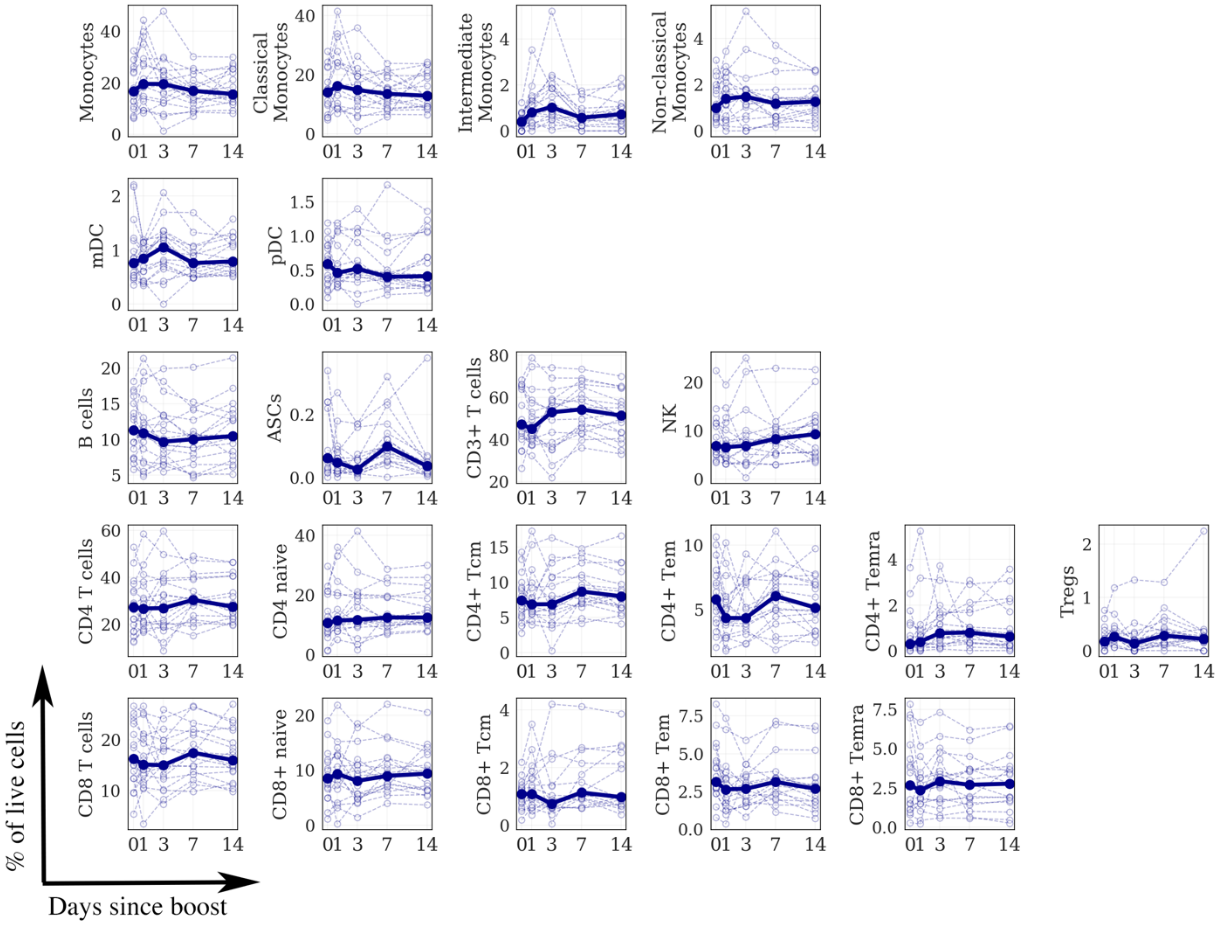


***Fig. S4. Longitudinal patterns of cell populations frequency profiled with CyTOF.*** Each graphic represents the longitudinal kinetics of the percentage of live cells from total PBMC for each individual population (y axis) determined by high-dimensional automated gated analysis of data generated by CyTOF assay. Data are expressed as connected time points from day 0 to day 1, 3, 7 and 14 post boost for each individual donor (thin lines) or the median of all (bold line).

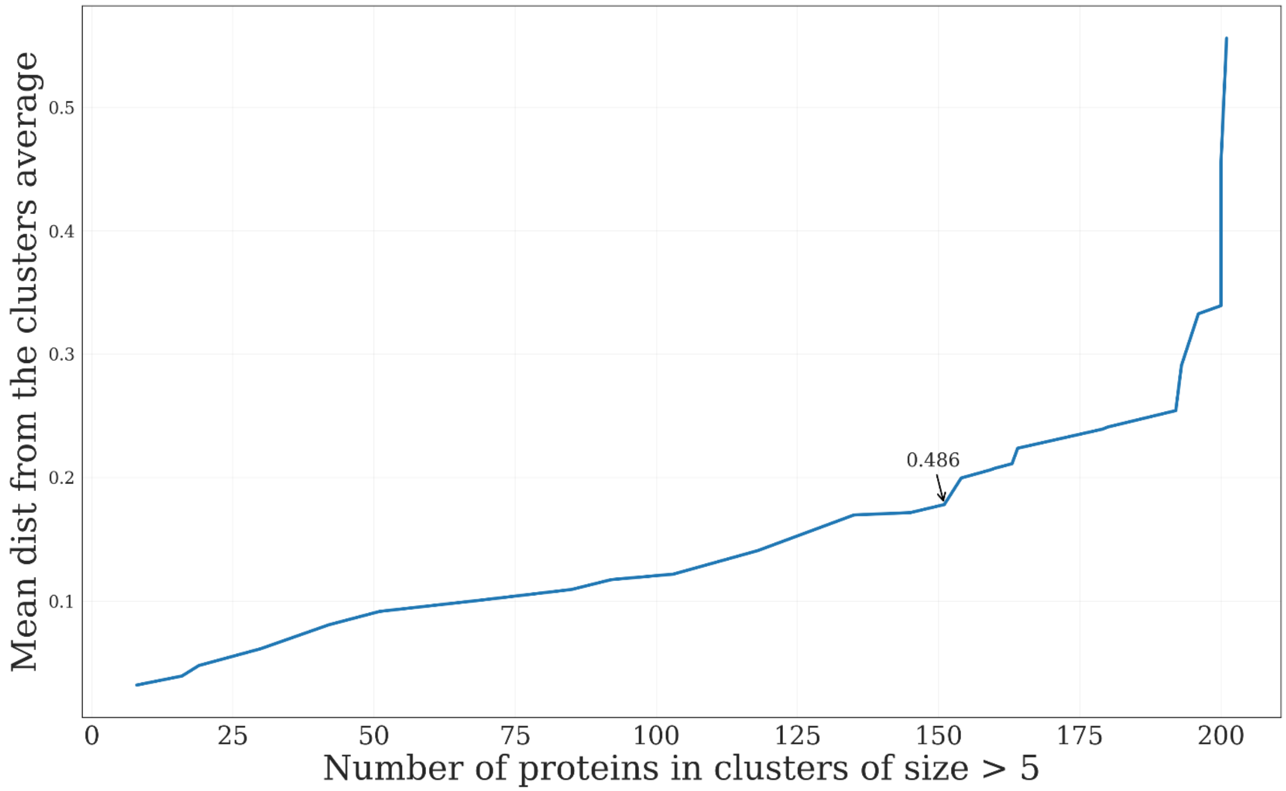


***Fig. S5. Cut-off threshold for proteomics data clustering.*** X-axis indicates number of proteins in clusters of size greater than 5. Y-axis indicates measure of cluster tightness. More specifically, it is correlation of every protein in the cluster with the cluster's mean, averaging across all the proteins in clusters containing 5 or more genes. Annotated point indicates a threshold of 0.486 chosen for downstream analysis. It indicates a fraction of maximum inter-protein distance.

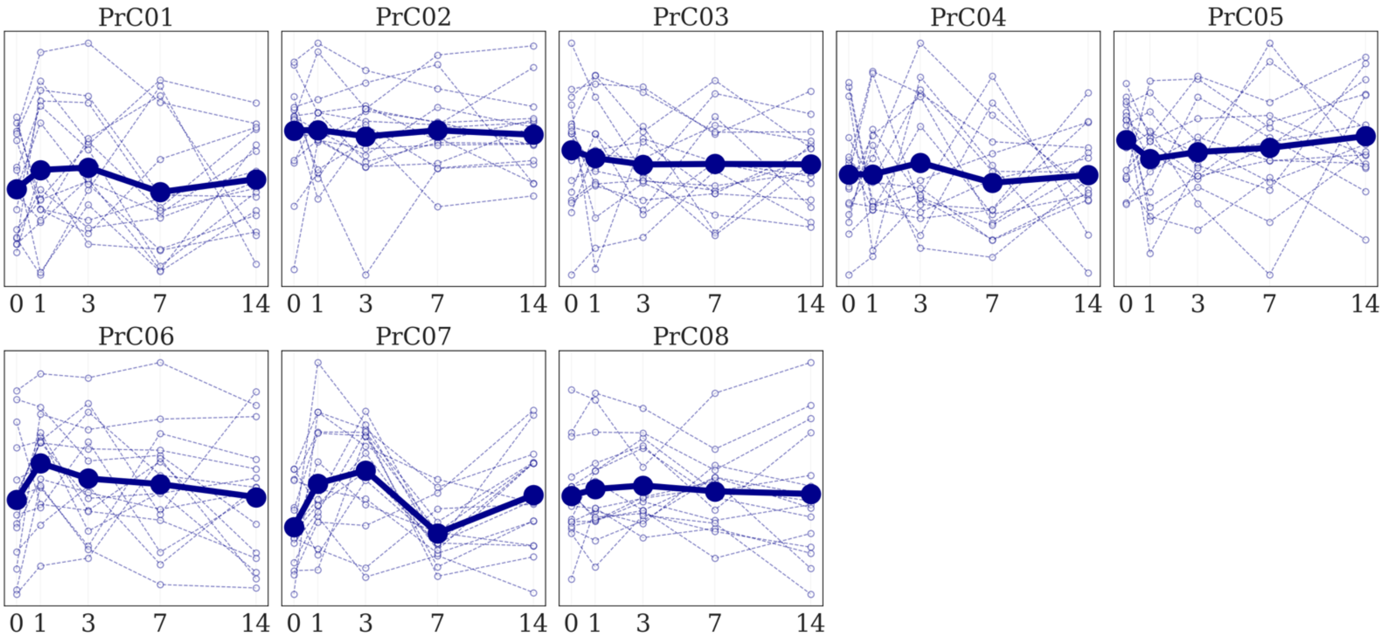


***Fig. S6*. *Longitudinal patterns of serum protein clusters.*** Thick lines represent average of PC1 for a given cluster. Averaging is done across donors of the same cohort: either primed with aP or wP vaccine. Thin lines in the background represent PC1 for every individual. Number of proteins comprising a cluster is given in brackets in the title above every plot.

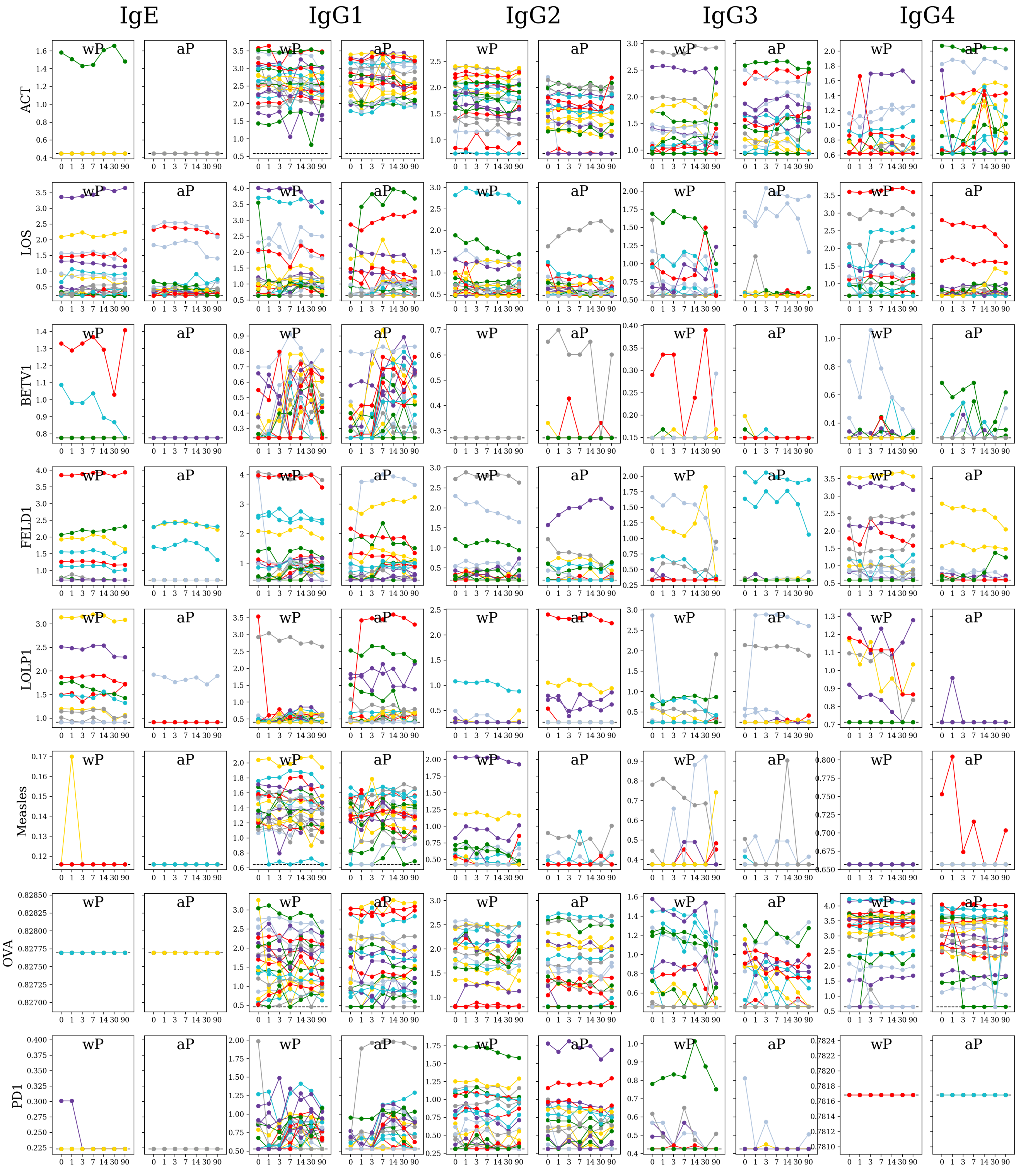


***Fig. S7. Quantification of immunoglobulins against whole-cell vaccine antigens, allergens and control antigens.*** Profiles of IgE and IgG1-4 versus the following antigens are shown: ACT (Adenylate Cyclase, wP antigen), LOS (lipopolysaccharide, wP antigen), BETV1 (seasonal allergen), FelD1 (allergen), LOLP1 (seasonal allergen), Measles (control), OVA (control), PD1 (control). Every immunoglobulin-antigen combination has two subplots, individually for DTwP- and DTaP-primed cohorts. Data is clipped to limit of detection. Y-axis indicates log10 of the IgE, IgG1, IgG2, IgG3, or IgG4 MFI. X-axis indicates days after vaccine boost (zero means pre-vaccination). There is not consistent induction of immunoglobulin levels at day 7 against any of these antigens unlike in the previous figure.

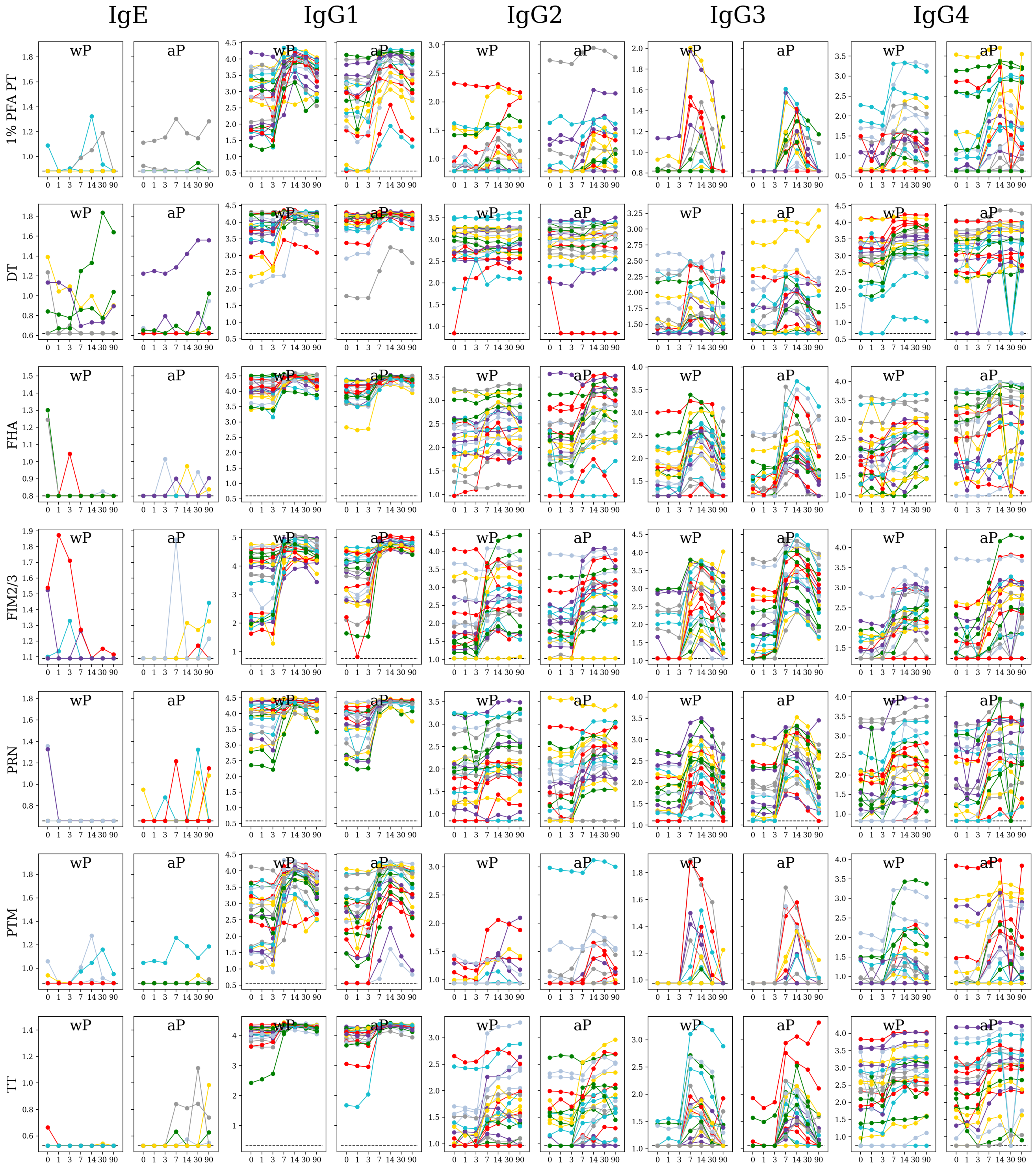


***Fig. S8. Quantification of Tdap antigen-specific immunoglobulins.*** Profiles of IgE and IgG1-4 in seven antigens from Tdap vaccine: 1% PFA PT (pertussis toxoid denaturized in 1% paraformaldehyde), DT (diphtheria toxoid), FHA (Filamentous hemagglutinin), FIM2/3 (Fimbriae 2/3), PRN (pertactin), PTM (pertussis toxin mutant), TT (tetanus toxoid). Every immunoglobulin-antigen combination has two subplots, individually for DTwP- and DTaP-primed cohorts. Note an increase of IgG immunoglobulins against all the antigens at day 7. Y-axis indicates log10 of the IgE, IgG1, IgG2, IgG3, or IgG4 MFI. X-axis indicates days after vaccine boost (zero means pre-vaccination).

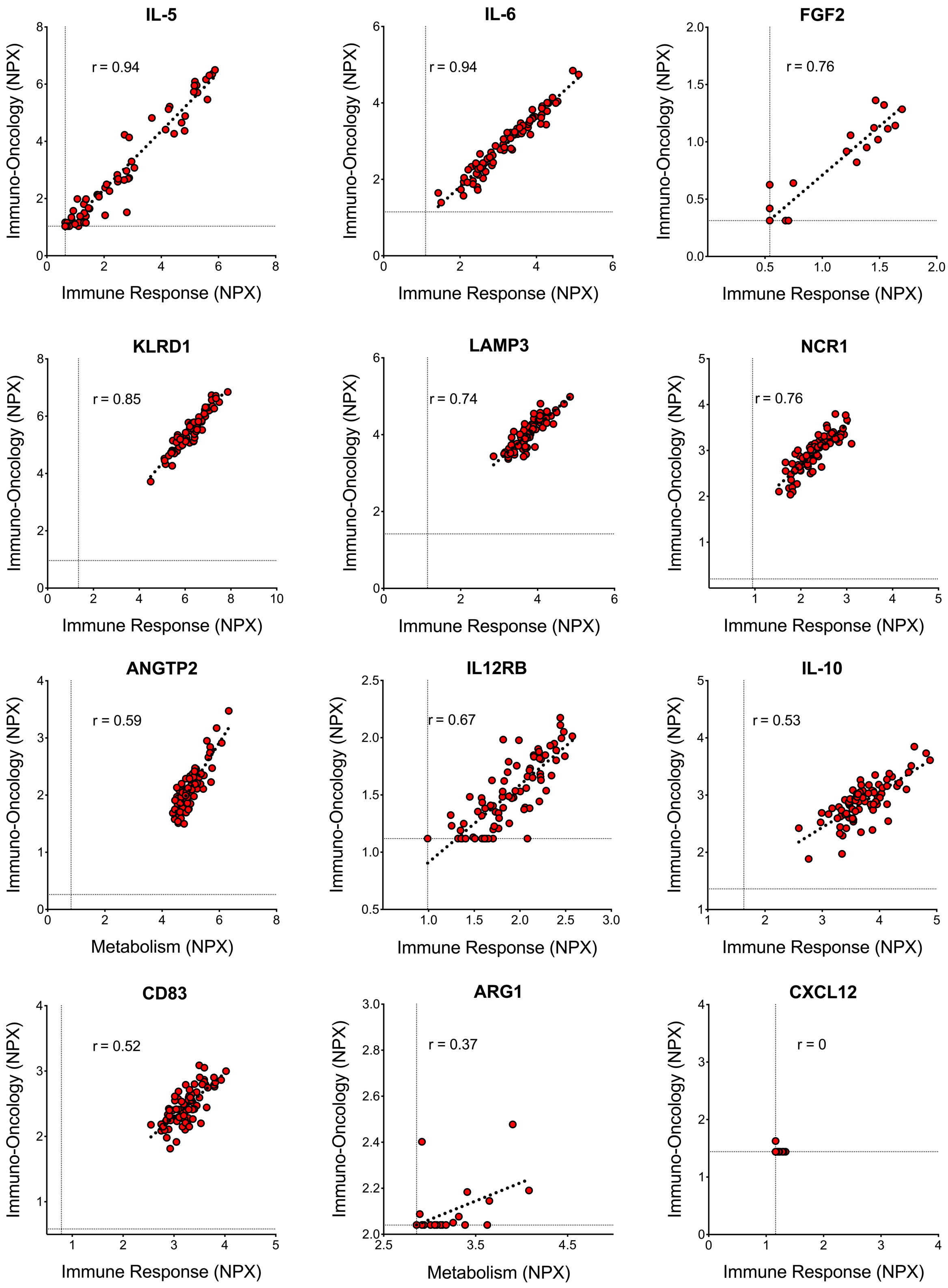


***Fig. S9. Correlation of duplicate analytes included in the proteomics analysis.*** Each axis contains the normalized protein expression values obtained by commercially available panels used to evaluate their presence in human plasma. Dotted line represents the limit of detection (LOD) of each analyte on each proximity extension assay panel (PEA). Correlation evaluation of the values obtained on each panel was performed with Spearman correlation test and r values for each individual analyte are represented on each graph. Values of r lower than 0.52 are obtained in analytes with the majority of NPX values equal or close to the LOD as observed in ARG1 and CXCL12.
