## Supplementary tables for "A system-view of *B. pertussis* booster vaccine responses in adults primed with whole-cell vs. acellular vaccine in infancy"

### Table S1 – Breakdown of assays performed per donor

| | | | Assays | | | | | |
|---|---|---|---|---|---|---|---|---|
| COUNT | DONOR ID | GROUP | SERUM IMUNOGLOBULINS | RNAseq | PLASMA PROTEOMICS | CyTOF | PLASMA ELISA | #Assays |
| 1 | 2730 | aP | TRUE | TRUE | TRUE | TRUE | TRUE | 4 |
| 2 | 2886 | aP | TRUE | TRUE | TRUE | TRUE | TRUE | 4 |
| 3 | 2902 | aP | TRUE | TRUE | TRUE | TRUE | TRUE | 4 |
| 4 | 2903 | aP | TRUE | TRUE | TRUE | TRUE | TRUE | 4 |
| 5 | 2904 | aP | TRUE | TRUE | TRUE | TRUE | TRUE | 4 |
| 6 | 2912 | aP | TRUE | TRUE | TRUE | TRUE | TRUE | 4 |
| 7 | 2915 | aP | TRUE | TRUE | TRUE | TRUE | TRUE | 4 |
| 8 | 2928 | aP | TRUE | TRUE | TRUE | TRUE | TRUE | 4 |
| 9 | 2685 | aP | TRUE | TRUE | FALSE | FALSE | TRUE | 2 |
| 10 | 2698 | aP | TRUE | TRUE | FALSE | FALSE | TRUE | 2 |
| 11 | 2728 | aP | TRUE | TRUE | FALSE | FALSE | TRUE | 2 |
| 12 | 2736 | aP | TRUE | TRUE | FALSE | FALSE | TRUE | 2 |
| 13 | 2900 | aP | TRUE | TRUE | FALSE | FALSE | TRUE | 2 |
| 14 | 2901 | aP | TRUE | TRUE | FALSE | FALSE | TRUE | 2 |
| 15 | 2923 | aP | TRUE | TRUE | FALSE | FALSE | TRUE | 2 |
| 16 | 2935 | aP | TRUE | TRUE | FALSE | FALSE | TRUE | 2 |
| 17 | 2689 | aP | TRUE | FALSE | FALSE | FALSE | FALSE | 1 |
| 18 | 2887 | aP | TRUE | FALSE | FALSE | FALSE | FALSE | 1 |
| 19 | 2890 | aP | TRUE | FALSE | FALSE | FALSE | FALSE | 1 |
| 20 | 2921 | aP | TRUE | FALSE | FALSE | FALSE | TRUE | 1 |
| 21 | 2922 | aP | TRUE | FALSE | FALSE | FALSE | FALSE | 1 |
| 22 | 2952 | aP | TRUE | FALSE | FALSE | FALSE | FALSE | 1 |
| 23 | 2968 | aP | TRUE | FALSE | FALSE | FALSE | TRUE | 1 |
| 24 | 2976 | aP | TRUE | FALSE | FALSE | FALSE | FALSE | 1 |
| 25 | 2995 | aP | TRUE | FALSE | FALSE | FALSE | FALSE | 1 |
| 26 | 2997 | aP | TRUE | FALSE | FALSE | FALSE | TRUE | 1 |
| 27 | 2998 | aP | TRUE | FALSE | FALSE | FALSE | TRUE | 1 |
| 28 | 3000 | aP | TRUE | FALSE | FALSE | FALSE | TRUE | 1 |
| 29 | 1829 | wP | TRUE | TRUE | TRUE | TRUE | TRUE | 4 |
| 30 | 2383 | wP | TRUE | TRUE | TRUE | TRUE | TRUE | 4 |
| 31 | 2687 | wP | TRUE | TRUE | TRUE | TRUE | TRUE | 4 |
| 32 | 2691 | wP | TRUE | TRUE | TRUE | TRUE | TRUE | 4 |
| 33 | 2696 | wP | TRUE | TRUE | TRUE | TRUE | TRUE | 4 |
| 34 | 2702 | wP | TRUE | TRUE | TRUE | TRUE | TRUE | 4 |
| 35 | 2703 | wP | TRUE | TRUE | TRUE | TRUE | TRUE | 4 |
| 36 | 2726 | wP | TRUE | TRUE | TRUE | TRUE | TRUE | 4 |
| 37 | 2732 | wP | TRUE | TRUE | TRUE | TRUE | TRUE | 4 |
| 38 | 2738 | wP | TRUE | TRUE | TRUE | TRUE | TRUE | 4 |
| 39 | 1655 | wP | TRUE | TRUE | FALSE | FALSE | TRUE | 2 |
| 40 | 1687 | wP | TRUE | TRUE | FALSE | FALSE | FALSE | 2 |
| 41 | 1832 | wP | TRUE | TRUE | FALSE | FALSE | TRUE | 2 |
| 42 | 2686 | wP | TRUE | TRUE | FALSE | FALSE | FALSE | 2 |
| 43 | 2699 | wP | TRUE | TRUE | FALSE | FALSE | FALSE | 2 |
| 44 | 2704 | wP | TRUE | TRUE | FALSE | FALSE | TRUE | 2 |
| 45 | 2706 | wP | TRUE | TRUE | FALSE | FALSE | TRUE | 2 |
| 46 | 2707 | wP | TRUE | TRUE | FALSE | FALSE | TRUE | 2 |
| 47 | 2719 | wP | TRUE | TRUE | FALSE | FALSE | TRUE | 2 |
| 48 | 2884 | wP | TRUE | TRUE | FALSE | FALSE | TRUE | 2 |
| 49 | 2500 | wP | TRUE | FALSE | FALSE | FALSE | FALSE | 1 |
| 50 | 2688 | wP | TRUE | FALSE | FALSE | FALSE | FALSE | 1 |
| 51 | 2690 | wP | TRUE | FALSE | FALSE | FALSE | FALSE | 1 |
| 52 | 2693 | wP | TRUE | FALSE | FALSE | FALSE | FALSE | 1 |
| 53 | 2729 | wP | TRUE | FALSE | FALSE | FALSE | FALSE | 1 |
| 54 | 2731 | wP | TRUE | FALSE | FALSE | FALSE | FALSE | 1 |
| 55 | 2883 | wP | TRUE | FALSE | FALSE | FALSE | FALSE | 1 |
| 56 | 2893 | wP | TRUE | FALSE | FALSE | FALSE | FALSE | 1 |
| 57 | 2895 | wP | TRUE | FALSE | FALSE | FALSE | FALSE | 1 |
| 58 | 2899 | wP | TRUE | FALSE | FALSE | FALSE | FALSE | 1 |
| Total donors per assay | | | 58 | 36 | 18 | 18 | 38 | 58 |

Table S2 – Transcriptomic clusters perturbation Wilcoxon p-values

| | Wilcoxon p-value | | |
|---|---|---|---|
| | all | wP | aP |
| TrC01 | 6.72E-04 | 2.27E-02 | 3.36E-03 |
| TrC02 | 1.10E-01 | 3.30E-01 | 7.71E-02 |
| TrC03 | 5.52E-05 | 1.12E-02 | 3.36E-03 |
| TrC04 | 1.03E-03 | 3.82E-02 | 1.13E-02 |
| TrC05 | 1.53E-04 | 7.78E-03 | 1.13E-02 |
| TrC06 | 1.09E-02 | 5.83E-02 | 4.86E-02 |
| TrC07 | 1.81E-02 | 1.79E-02 | 1.27E-01 |
| TrC08 | 2.91E-02 | 1.62E-01 | 8.73E-02 |
| TrC09 | 1.42E-04 | 1.34E-02 | 3.36E-03 |
| TrC10 | 8.69E-06 | 2.54E-03 | 3.36E-03 |
| TrC11 | 6.65E-04 | 1.52E-02 | 3.10E-02 |
| TrC12 | 2.32E-01 | 7.96E-02 | 6.05E-01 |
| TrC13 | 8.20E-05 | 5.73E-03 | 7.54E-03 |
| TrC14 | 3.89E-01 | 3.91E-01 | 1.98E-01 |
| TrC15 | 1.09E-02 | 9.10E-02 | 8.05E-02 |
| TrC16 | 7.20E-05 | 4.90E-03 | 1.09E-02 |
| TrC17 | 5.48E-06 | 1.09E-03 | 5.30E-02 |
| TrC18 | 6.69E-03 | 4.70E-03 | 1.37E-01 |
| TrC19 | 3.65E-01 | 2.68E-01 | 3.45E-01 |
| TrC20 | 5.42E-03 | 4.41E-03 | 5.29E-01 |
| TrC21 | 4.37E-02 | 4.41E-03 | 8.05E-02 |
| TrC22 | 7.55E-03 | 1.74E-02 | 4.55E-01 |
| TrC23 | 8.42E-03 | 5.42E-03 | 5.49E-01 |
| TrC24 | 5.10E-03 | 2.43E-02 | 4.46E-02 |
| TrC25 | 1.15E-04 | 1.62E-02 | 5.25E-03 |
| TrC26 | 1.88E-04 | 1.79E-02 | 6.85E-03 |
| TrC27 | 5.48E-06 | 1.09E-03 | 3.36E-03 |
| TrC28 | 1.42E-04 | 1.91E-02 | 3.36E-03 |
| TrC29 | 5.48E-06 | 6.47E-03 | 3.36E-03 |
| TrC30 | 3.04E-05 | 1.09E-03 | 3.36E-03 |
| TrC31 | 2.32E-05 | 1.09E-03 | 8.33E-03 |
| TrC32 | 7.55E-03 | 9.34E-03 | 4.74E-01 |
| TrC33 | 4.99E-01 | 3.60E-01 | 2.32E-01 |
| TrC34 | 2.95E-05 | 4.53E-03 | 6.24E-03 |
| TrC35 | 3.67E-05 | 6.08E-03 | 6.24E-03 |
| TrC36 | 9.36E-05 | 5.14E-03 | 7.71E-02 |
| TrC37 | 1.39E-03 | 1.52E-02 | 9.46E-02 |
| TrC38 | 1.47E-05 | 1.09E-03 | 7.42E-02 |
| TrC39 | 8.69E-06 | 3.31E-03 | 3.36E-03 |

### Table S3 – Cell frequencies vs transcriptomic cluster Spearman correlation r values

| Cluster ID | Monocytes | CD33+HLADR+ | Classical_Monocytes | Non-Classical_Monocytes | Intermediate_Monocytes | Bcells | CD3CD19 | CD3CD19neg | CD3 Tcells | CD4Tcells | CD8Tcells | Tregs | TemraCD4 | NaiveCD4 | TemCD4 | TcmCD4 | TemraCD8 | NaiveCD8 | TemCD8 | TcmCD8 | NK | Basophils | mDC | pDC |
|---|---|---|---|---|---|---|---|---|---|---|---|---|---|---|---|---|---|---|---|---|---|---|---|---|
| **TrC01** | -0.14 | -0.14 | -0.17 | -0.01 | 0.12 | -0.21 | -0.02 | 0.09 | 0.09 | 0.11 | -0.02 | -0.08 | 0.19 | 0.06 | 0.09 | 0.17 | 0.14 | -0.09 | 0.09 | 0.16 | -0.15 | 0.07 | 0.10 | 0.07 |
| **TrC02** | -0.12 | -0.12 | -0.16 | -0.08 | 0.19 | -0.32 | 0.06 | 0.16 | -0.02 | -0.13 | 0.02 | -0.07 | 0.04 | -0.21 | -0.03 | 0.13 | 0.12 | -0.03 | 0.14 | 0.30 | -0.11 | 0.01 | 0.12 | -0.04 |
| **TrC03** | 0.13 | 0.13 | 0.16 | 0.00 | -0.09 | 0.21 | 0.05 | -0.11 | -0.07 | -0.07 | 0.07 | 0.13 | -0.19 | -0.09 | -0.03 | -0.06 | -0.09 | 0.09 | -0.03 | -0.03 | 0.15 | -0.06 | -0.13 | -0.12 |
| **TrC04** | -0.10 | -0.10 | -0.07 | -0.07 | -0.28 | 0.27 | 0.07 | -0.08 | 0.07 | 0.10 | 0.07 | 0.00 | -0.06 | 0.19 | -0.01 | -0.13 | -0.18 | 0.16 | -0.15 | -0.25 | 0.18 | -0.11 | -0.24 | -0.05 |
| **TrC05** | -0.20 | -0.20 | -0.17 | -0.20 | -0.22 | 0.13 | 0.18 | -0.08 | 0.10 | 0.09 | 0.24 | 0.01 | -0.17 | 0.10 | -0.03 | 0.04 | -0.19 | 0.30 | -0.07 | 0.06 | 0.15 | -0.11 | -0.28 | -0.16 |
| **TrC06** | 0.28 | 0.28 | 0.32 | 0.17 | -0.07 | 0.27 | -0.08 | -0.12 | -0.09 | -0.07 | -0.09 | 0.13 | -0.11 | -0.01 | -0.04 | -0.20 | -0.07 | -0.06 | -0.08 | -0.30 | 0.14 | -0.01 | 0.01 | 0.03 |
| **TrC07** | 0.54 | 0.54 | 0.56 | 0.20 | 0.16 | 0.20 | -0.11 | -0.15 | -0.30 | -0.28 | -0.12 | 0.18 | -0.30 | -0.32 | -0.08 | -0.16 | -0.01 | -0.13 | -0.02 | -0.09 | 0.04 | 0.07 | 0.15 | 0.03 |
| **TrC08** | 0.05 | 0.05 | 0.08 | 0.09 | -0.20 | 0.27 | -0.04 | -0.10 | 0.04 | 0.08 | -0.06 | 0.02 | 0.03 | 0.19 | -0.01 | -0.20 | -0.08 | 0.00 | -0.13 | -0.37 | 0.13 | -0.08 | -0.08 | 0.09 |
| **TrC09** | -0.09 | -0.10 | -0.13 | 0.05 | 0.08 | -0.14 | -0.06 | 0.08 | 0.09 | 0.14 | -0.08 | -0.08 | 0.26 | 0.14 | 0.10 | 0.09 | 0.15 | -0.14 | 0.07 | -0.01 | -0.15 | 0.04 | 0.10 | 0.10 |
| **TrC10** | -0.02 | -0.03 | -0.05 | 0.10 | 0.07 | -0.07 | -0.09 | 0.07 | 0.03 | 0.11 | -0.17 | -0.10 | 0.26 | 0.15 | 0.07 | -0.01 | 0.11 | -0.19 | -0.01 | -0.13 | -0.14 | 0.06 | 0.13 | 0.14 |
| **TrC11** | 0.74 | 0.74 | 0.76 | 0.30 | 0.28 | 0.20 | -0.17 | -0.12 | -0.43 | -0.40 | -0.28 | 0.15 | -0.31 | -0.46 | -0.09 | -0.22 | 0.05 | -0.29 | -0.06 | -0.13 | 0.03 | 0.12 | 0.33 | 0.14 |
| **TrC12** | 0.69 | 0.69 | 0.72 | 0.35 | 0.20 | 0.24 | -0.20 | -0.13 | -0.34 | -0.30 | -0.31 | 0.16 | -0.20 | -0.30 | -0.09 | -0.27 | 0.05 | -0.31 | -0.10 | -0.30 | 0.06 | 0.10 | 0.31 | 0.19 |
| **TrC13** | -0.19 | -0.18 | -0.18 | -0.22 | -0.05 | -0.04 | 0.25 | -0.01 | 0.02 | -0.10 | 0.26 | 0.00 | -0.25 | -0.13 | -0.13 | 0.07 | -0.12 | 0.29 | 0.01 | 0.30 | 0.09 | -0.09 | -0.20 | -0.20 |
| **TrC14** | 0.57 | 0.56 | 0.59 | 0.31 | 0.11 | 0.20 | -0.17 | -0.12 | -0.26 | -0.24 | -0.27 | 0.17 | -0.14 | -0.21 | -0.06 | -0.26 | 0.03 | -0.26 | -0.09 | -0.34 | 0.08 | 0.03 | 0.26 | 0.17 |
| **TrC15** | -0.13 | -0.12 | -0.15 | -0.11 | 0.17 | -0.33 | 0.12 | 0.14 | -0.04 | -0.18 | 0.06 | 0.01 | -0.01 | -0.28 | -0.06 | 0.12 | 0.09 | 0.00 | 0.13 | 0.36 | -0.08 | -0.01 | 0.03 | -0.13 |
| **TrC16** | -0.03 | -0.02 | -0.04 | -0.09 | 0.16 | -0.22 | 0.15 | 0.10 | -0.11 | -0.28 | 0.10 | 0.05 | -0.18 | -0.39 | -0.12 | 0.07 | 0.02 | 0.07 | 0.12 | 0.36 | 0.00 | -0.06 | 0.00 | -0.16 |
| **TrC17** | 0.70 | 0.71 | 0.72 | 0.44 | 0.30 | 0.21 | -0.24 | -0.04 | -0.40 | -0.33 | -0.40 | 0.04 | -0.10 | -0.34 | -0.07 | -0.29 | 0.13 | -0.41 | -0.08 | -0.35 | -0.02 | 0.10 | 0.39 | 0.26 |
| **TrC18** | -0.32 | -0.34 | -0.31 | -0.12 | -0.30 | 0.20 | 0.09 | -0.05 | 0.23 | 0.30 | 0.09 | -0.11 | 0.12 | 0.46 | 0.01 | -0.08 | -0.19 | 0.19 | -0.16 | -0.25 | 0.11 | -0.10 | -0.26 | 0.02 |
| **TrC19** | 0.20 | 0.20 | 0.22 | 0.22 | -0.09 | 0.29 | -0.08 | -0.09 | -0.04 | 0.02 | -0.16 | 0.04 | 0.06 | 0.12 | -0.01 | -0.22 | -0.01 | -0.13 | -0.12 | -0.41 | 0.09 | -0.06 | 0.02 | 0.12 |
| **TrC20** | -0.54 | -0.56 | -0.53 | -0.35 | -0.39 | -0.02 | 0.06 | -0.14 | 0.43 | 0.58 | 0.24 | -0.06 | 0.14 | 0.66 | 0.17 | 0.24 | -0.19 | 0.31 | -0.11 | 0.05 | -0.09 | 0.02 | -0.27 | 0.02 |
| **TrC21** | 0.05 | 0.05 | 0.04 | -0.06 | 0.04 | -0.29 | -0.14 | 0.10 | -0.11 | 0.00 | -0.12 | 0.01 | 0.09 | -0.06 | 0.17 | 0.02 | -0.25 | -0.08 | 0.01 | 0.02 | -0.10 | 0.22 | 0.13 | 0.13 |
| **TrC22** | -0.19 | -0.15 | -0.22 | 0.23 | 0.04 | 0.07 | 0.17 | 0.70 | -0.28 | -0.36 | -0.14 | -0.41 | 0.09 | -0.28 | -0.03 | -0.43 | 0.11 | -0.16 | 0.12 | -0.36 | 0.63 | -0.47 | -0.13 | -0.20 |
| **TrC23** | 0.67 | 0.67 | 0.68 | 0.25 | 0.40 | 0.00 | -0.13 | -0.02 | -0.49 | -0.50 | -0.28 | 0.12 | -0.30 | -0.62 | -0.11 | -0.15 | 0.07 | -0.29 | -0.03 | 0.08 | -0.04 | 0.12 | 0.34 | 0.08 |
| **TrC24** | -0.34 | -0.32 | -0.33 | 0.02 | -0.08 | 0.74 | 0.36 | 0.16 | -0.13 | -0.04 | 0.05 | -0.28 | 0.00 | 0.09 | -0.22 | -0.18 | -0.03 | 0.15 | -0.29 | -0.04 | 0.24 | -0.31 | -0.45 | -0.31 |
| **TrC25** | 0.32 | 0.33 | 0.36 | 0.20 | -0.08 | 0.27 | -0.16 | -0.11 | -0.13 | -0.06 | -0.12 | 0.19 | -0.10 | -0.03 | 0.07 | -0.16 | -0.02 | -0.13 | -0.02 | -0.26 | 0.13 | 0.01 | 0.02 | 0.02 |
| **TrC26** | 0.07 | 0.07 | 0.08 | 0.18 | -0.04 | 0.22 | -0.06 | -0.04 | 0.00 | 0.07 | -0.17 | -0.09 | 0.13 | 0.21 | -0.01 | -0.21 | 0.04 | -0.14 | -0.10 | -0.35 | -0.01 | -0.06 | 0.04 | 0.11 |
| **TrC27** | -0.09 | -0.11 | -0.06 | -0.06 | -0.24 | 0.30 | 0.05 | -0.18 | 0.20 | 0.21 | 0.16 | 0.09 | 0.09 | 0.27 | -0.03 | 0.03 | -0.19 | 0.27 | -0.01 | -0.10 | 0.07 | 0.02 | -0.31 | -0.19 |
| **TrC28** | 0.37 | 0.38 | 0.38 | 0.12 | 0.21 | -0.01 | 0.00 | -0.03 | -0.30 | -0.37 | -0.07 | 0.17 | -0.25 | -0.50 | -0.07 | -0.03 | 0.06 | -0.13 | 0.07 | 0.19 | 0.04 | -0.02 | 0.11 | -0.09 |
| **TrC29** | 0.36 | 0.36 | 0.36 | 0.34 | 0.17 | 0.15 | -0.16 | 0.01 | -0.20 | -0.08 | -0.36 | -0.04 | 0.15 | -0.04 | 0.01 | -0.20 | 0.11 | -0.35 | -0.09 | -0.37 | -0.05 | 0.02 | 0.22 | 0.20 |
| **TrC30** | 0.34 | 0.34 | 0.34 | 0.25 | 0.18 | 0.08 | -0.19 | -0.08 | -0.12 | 0.02 | -0.26 | 0.01 | 0.19 | -0.03 | 0.10 | 0.00 | 0.17 | -0.31 | 0.01 | -0.14 | -0.22 | 0.11 | 0.19 | 0.11 |
| **TrC31** | 0.20 | 0.18 | 0.20 | 0.02 | 0.12 | 0.22 | -0.07 | -0.08 | -0.12 | -0.03 | -0.14 | -0.18 | -0.08 | 0.03 | -0.15 | -0.17 | 0.09 | -0.10 | -0.24 | -0.25 | -0.10 | 0.10 | 0.09 | 0.32 |
| **TrC32** | -0.63 | -0.66 | -0.63 | -0.52 | -0.38 | -0.30 | 0.04 | -0.14 | 0.50 | 0.61 | 0.38 | -0.01 | 0.08 | 0.61 | 0.24 | 0.44 | -0.23 | 0.45 | 0.00 | 0.33 | -0.16 | 0.09 | -0.29 | -0.01 |
| **TrC33** | -0.09 | -0.10 | -0.05 | -0.18 | -0.08 | 0.15 | 0.07 | -0.14 | 0.04 | 0.20 | 0.13 | -0.06 | 0.08 | 0.14 | 0.20 | 0.11 | 0.10 | 0.11 | 0.14 | 0.12 | -0.21 | 0.04 | -0.26 | -0.25 |
| **TrC34** | -0.21 | -0.20 | -0.18 | -0.17 | -0.20 | -0.02 | 0.08 | 0.04 | 0.06 | 0.07 | 0.14 | 0.14 | 0.03 | 0.00 | 0.05 | 0.12 | -0.13 | 0.16 | 0.00 | 0.14 | 0.17 | -0.12 | -0.25 | -0.22 |
| **TrC35** | -0.13 | -0.13 | -0.13 | 0.06 | -0.08 | 0.03 | -0.04 | 0.03 | 0.12 | 0.15 | -0.10 | -0.14 | 0.22 | 0.30 | 0.01 | -0.15 | -0.06 | -0.04 | -0.08 | -0.32 | -0.04 | 0.04 | 0.01 | 0.16 |
| **TrC36** | -0.64 | -0.65 | -0.62 | -0.54 | -0.36 | -0.12 | 0.19 | -0.10 | 0.39 | 0.46 | 0.39 | 0.02 | 0.02 | 0.52 | 0.09 | 0.30 | -0.27 | 0.51 | -0.14 | 0.32 | -0.03 | 0.03 | -0.37 | -0.13 |
| **TrC37** | 0.11 | 0.11 | 0.12 | 0.17 | -0.03 | 0.03 | -0.12 | -0.14 | 0.15 | 0.16 | 0.10 | 0.21 | 0.09 | 0.10 | 0.13 | 0.22 | 0.08 | 0.00 | 0.25 | 0.17 | -0.09 | 0.11 | -0.02 | -0.24 |
| **TrC38** | 0.43 | 0.41 | 0.44 | 0.10 | 0.15 | 0.30 | -0.05 | -0.18 | -0.22 | -0.17 | -0.17 | -0.10 | -0.23 | -0.12 | -0.20 | -0.23 | 0.01 | -0.13 | -0.27 | -0.24 | -0.08 | 0.16 | 0.17 | 0.30 |
| **TrC39** | -0.18 | -0.18 | -0.17 | -0.17 | -0.12 | -0.14 | 0.08 | 0.06 | 0.04 | 0.00 | 0.12 | 0.20 | 0.05 | -0.10 | 0.06 | 0.18 | -0.06 | 0.11 | 0.07 | 0.26 | 0.09 | -0.09 | -0.19 | -0.23 |

Table S4 - Transcriptomic clusters Mann-Whitney p-values group comparison

| | Days post boost | | | | |
|---|---|---|---|---|---|
| **Cluster** | **0d** | **1d** | **3d** | **7d** | **14d** |
| **TrC01** | 0.06 | 0.21 | 0.59 | 0.22 | 0.12 |
| **TrC02** | 0.07 | 0.26 | 0.59 | 0.23 | 0.12 |
| **TrC03** | 0.06 | 0.34 | 0.59 | 0.30 | 0.27 |
| **TrC04** | 0.06 | 0.08 | 0.59 | 0.45 | 0.27 |
| **TrC05** | 0.30 | 0.57 | 0.59 | 0.78 | 0.57 |
| **TrC06** | 0.04 | 0.11 | 0.59 | 0.28 | 0.24 |
| **TrC07** | 0.14 | 0.72 | 0.75 | 0.30 | 0.27 |
| **TrC08** | 0.06 | 0.04 | 0.59 | 0.30 | 0.35 |
| **TrC09** | 0.33 | 0.34 | 0.59 | 0.23 | 0.13 |
| **TrC10** | 0.73 | 0.43 | 0.59 | 0.30 | 0.14 |
| **TrC11** | 0.47 | 0.96 | 0.99 | 0.30 | 0.40 |
| **TrC12** | 0.06 | 0.51 | 0.66 | 0.22 | 0.27 |
| **TrC13** | 0.91 | 0.96 | 0.90 | 0.86 | 0.96 |
| **TrC14** | 0.04 | 0.34 | 0.59 | 0.23 | 0.27 |
| **TrC15** | 0.33 | 0.28 | 0.59 | 0.30 | 0.21 |
| **TrC16** | 0.96 | 0.51 | 0.59 | 0.66 | 0.39 |
| **TrC17** | 0.14 | 0.96 | 0.93 | 0.22 | 0.54 |
| **TrC18** | 0.47 | 0.34 | 0.76 | 0.84 | 0.93 |
| **TrC19** | 0.07 | 0.08 | 0.59 | 0.23 | 0.27 |
| **TrC20** | 0.53 | 0.96 | 0.65 | 0.66 | 0.27 |
| **TrC21** | 0.53 | 0.66 | 0.90 | 0.23 | 0.90 |
| **TrC22** | 0.33 | 0.38 | 0.59 | 0.85 | 0.99 |
| **TrC23** | 0.82 | 0.34 | 0.59 | 0.86 | 0.99 |
| **TrC24** | 0.33 | 0.21 | 0.59 | 0.30 | 0.21 |
| **TrC25** | 0.06 | 0.28 | 0.59 | 0.23 | 0.27 |
| **TrC26** | 0.64 | 0.34 | 0.59 | 0.38 | 0.99 |
| **TrC27** | 0.00 | 0.02 | 0.21 | 0.23 | 0.04 |
| **TrC28** | 0.47 | 0.72 | 0.66 | 0.85 | 0.82 |
| **TrC29** | 0.33 | 0.96 | 0.59 | 0.30 | 0.50 |
| **TrC30** | 0.91 | 0.56 | 0.59 | 0.58 | 0.21 |
| **TrC31** | 0.91 | 0.34 | 0.38 | 0.01 | 0.27 |
| **TrC32** | 0.15 | 0.57 | 0.66 | 0.22 | 0.13 |
| **TrC33** | 0.96 | 0.96 | 0.70 | 0.30 | 0.65 |
| **TrC34** | 0.06 | 0.90 | 0.90 | 0.84 | 0.73 |
| **TrC35** | 0.82 | 0.96 | 0.99 | 0.86 | 0.27 |
| **TrC36** | 0.81 | 0.96 | 0.99 | 0.30 | 0.40 |
| **TrC37** | 0.10 | 0.36 | 0.59 | 0.84 | 0.51 |
| **TrC38** | 0.54 | 0.36 | 0.38 | 0.01 | 0.04 |
| **TrC39** | 0.14 | 0.69 | 0.59 | 0.85 | 0.41 |

Table S5 – Proteomics excluded assays and exclusion criteria Olink PEA

| Count | Olink panel | Assay | Uniprot ID | Reason for exclusion |
|---|---|---|---|---|
| 1 | Olink IMMUNE RESPONSE(v.3202) | ARNT | P27540 | ≤ LOD |
| 2 | Olink IMMUNE RESPONSE(v.3202) | BIRC2 | Q13490 | ≤ LOD |
| 3 | Olink IMMUNE RESPONSE(v.3202) | DGKZ | Q13574 | ≤ LOD |
| 4 | Olink IMMUNE RESPONSE(v.3202) | EIF5A | P63241 | ≤ LOD |
| 5 | Olink METABOLISM(v.3402) | GLRX | P35754 | ≤ LOD |
| 6 | Olink IMMUNO-ONCOLOGY(v.3101) | IFN-beta | P01574 | ≤ LOD |
| 7 | Olink IMMUNO-ONCOLOGY(v.3101) | IFN-gamma | P01579 | ≤ LOD |
| 8 | Olink IMMUNE RESPONSE(v.3202) | IFNLR1 | Q8IU57 | ≤ LOD |
| 9 | Olink IMMUNO-ONCOLOGY(v.3101) | IL-1 alpha | P01583 | ≤ LOD |
| 10 | Olink IMMUNO-ONCOLOGY(v.3101) | IL-21 | Q9HBE4 | ≤ LOD |
| 11 | Olink IMMUNO-ONCOLOGY(v.3101) | IL-35 | Q14213,P29459 | ≤ LOD |
| 12 | Olink IMMUNO-ONCOLOGY(v.3101) | IL13 | P35225 | ≤ LOD |
| 13 | Olink IMMUNO-ONCOLOGY(v.3101) | IL2 | P60568 | ≤ LOD |
| 14 | Olink IMMUNO-ONCOLOGY(v.3101) | IL33 | O95760 | ≤ LOD |
| 15 | Olink IMMUNE RESPONSE(v.3202) | IRAK4 | Q9NWZ3 | ≤ LOD |
| 16 | Olink IMMUNE RESPONSE(v.3202) | KPNA1 | P52294 | ≤ LOD |
| 17 | Olink IMMUNE RESPONSE(v.3202) | NF2 | P35240 | ≤ LOD |
| 18 | Olink METABOLISM(v.3402) | NQO2 | P16083 | ≤ LOD |
| 19 | Olink IMMUNO-ONCOLOGY(v.3101) | PTN | P21246 | ≤ LOD |
| 20 | Olink METABOLISM(v.3402) | S100P | P25815 | ≤ LOD |
| 21 | Olink IMMUNO-ONCOLOGY(v.3101) | TNF | P01375 | ≤ LOD |
| 22 | Olink IMMUNE RESPONSE(v.3202) | CD28 | P10747 | ≤ LOD |
| 23 | Olink IMMUNO-ONCOLOGY(v.3101) | CD28 | P10747 | ≤ LOD |
| 24 | Olink IMMUNO-ONCOLOGY(v.3101) | CD83 | Q01151 | Duplicate |
| 25 | Olink IMMUNO-ONCOLOGY(v.3101) | CXCL12 | P48061 | Duplicate |
| 26 | Olink IMMUNO-ONCOLOGY(v.3101) | FGF2 | P09038 | Duplicate |
| 27 | Olink IMMUNO-ONCOLOGY(v.3101) | IL10 | P22301 | Duplicate |
| 28 | Olink IMMUNO-ONCOLOGY(v.3101) | IL12RB1 | P42701 | Duplicate |
| 29 | Olink IMMUNO-ONCOLOGY(v.3101) | IL5 | P05113 | Duplicate |
| 30 | Olink IMMUNO-ONCOLOGY(v.3101) | IL6 | P05231 | Duplicate |
| 31 | Olink IMMUNO-ONCOLOGY(v.3101) | KLRD1 | Q13241 | Duplicate |
| 32 | Olink IMMUNO-ONCOLOGY(v.3101) | LAMP3 | Q9UQV4 | Duplicate |
| 33 | Olink IMMUNO-ONCOLOGY(v.3101) | NCR1 | O76036 | Duplicate |
| 34 | Olink IMMUNO-ONCOLOGY(v.3101) | ANGPT2 | O15123 | Duplicate |
| 35 | Olink IMMUNO-ONCOLOGY(v.3101) | ARG1 | P05089 | Duplicate |

Table S6 – CyTOF Antibodies and conjugations

| Marker | Metal | Clone |
|---|---|---|
| CD45 | 89Y | HI30 |
| CD3 | 115In | UCHT1 |
| CD19 | 142Nd | HIB19 |
| CD38 | 144Nd | HIT2 |
| CD4 | 145Nd | RPA-T4 |
| CD20 | 145Nd | 2H7 |
| CD123 | 151Eu | 6H6 |
| CD45RA | 155Gd | HI100 |
| CD1c | 160Gd | L161 |
| CD33 | 163Dy | WM53 |
| CCR7 | 167Er | G043H7 |
| CD25 | 169Tm | M-A251 |
| CD8a | 172Yb | RPA-T8 |
| CD14 | 173Yb | 61D3 |
| HLA-DR | 174Yb | L243 |
| CD56 | 176Yb | CMSSB |
| CD16 | 209Bi | 3G8 |
